# Phagocytosis-driven neurodegeneration through opposing roles of an ABC transporter in neurons and phagocytes

**DOI:** 10.1101/2024.05.19.594870

**Authors:** Xinchen Chen, Bei Wang, Ankita Sarkar, Zixian Huang, Nicolas Vergara Ruiz, Ann T. Yeung, Rachael Chen, Chun Han

## Abstract

Lipid homeostasis is critical to the survival of neurons. Lipid transporters from the ATP-binding cassette A (ABCA) subfamily are important regulators of lipid trafficking and are associated with multiple neurodegenerative diseases. How ABCA transporters regulate specific aspects of lipid homeostasis to impact neurodegeneration is an outstanding question. Here we report that the *Drosophila* ABCA protein Engulfment ABC Transporter in the ovary (Eato) contributes to phagocytosis-dependent neurodegeneration by playing two opposing roles in neurons and nearby phagocytes: In neurons, Eato prevents dendrites and axons from being attacked and engulfed by neighboring phagocytes; in phagocytes, however, Eato enhances the ability of these cells to detect neurons as engulfment targets. Thus, *Eato* deficiency in neurons alone results in severe phagocytosis-dependent dendrite and axon degeneration, whereas removing *Eato* from both neurons and phagocytes completely rescues the neurite degeneration. Surprisingly, Eato exerts its functions in both neurons and phagocytes by suppressing the effects of the eat-me signal phosphatidylserine (PS) exposed on the cell surface. Interestingly, multiple human and *C. elegans* ABCA homologs can compensate for the loss of *Eato* in phagocytes but not in neurons, suggesting both conserved and cell type-specific activities of these ABCA proteins. These results reveal how ABCA proteins participate in neurodegeneration by regulating PS homeostasis and imply possible mechanisms of neuron-phagocyte interactions in neurodegenerative diseases.

## INTRODUCTION

For a neuron to survive and function, the composition and spatial distribution of its lipids need to be dynamically maintained within a narrow range of optimal levels. Lipid homeostasis is controlled largely by lipid transporters located on cell membranes. ATP-binding cassette (ABC) proteins comprise a large family of transporters that contain conserved nucleotide-binding domains (NBDs) and that translocate diverse substrates across cell membranes using energy from ATP (Dean et al., 2022; Quazi and Molday, 2011). Among ABC proteins, the ABCA subfamily is best known for its functions in transporting lipids across the bilayer and loading lipids onto apolipoprotein carriers (Albrecht and Viturro, 2007; Broccardo et al., 1999; Quazi and Molday, 2011). The importance of ABCA genes in human health is underscored by numerous mutations that are associated with diverse inheritable diseases related to lipid transport, including multiple neurodegenerative diseases (Albrecht and Viturro, 2007; Bossaerts et al., 2022; Piehler et al., 2012).

Several ABCA proteins in humans and other animals are protective of neurons. Genome-wide association studies identified *ABCA1* and *ABCA7* as risk genes for Alzheimer’s disease (AD) (Bellenguez et al., 2022; Hollingworth et al., 2011). In mice, ABCA1 promotes the efflux of excess cholesterols from the brain and the lipidation of apolipoproteins, including the AD-associated ApoE (Do et al., 2011; Kim et al., 2007; Wahrle et al., 2004). High membrane cholesterol promotes production of the neurotoxic amyloid beta peptide (Aβ) (Anstey et al., 2017; Cho et al., 2020), while lipidated ApoE can bind Aβ and is negatively correlated with AD (Tokuda et al., 2000; Zhao et al., 2017). ABCA7 reduces Aβ production by affecting amyloid precursor protein (APP) processing (Sakae et al., 2016; Satoh et al., 2015) and can also reduce the buildup of extracellular Aβ by promoting phagocytosis (Fu et al., 2016; Kim et al., 2013). Unlike most ABCA proteins, which act as floppases to export lipids from the cytosolic to the extracellular leaflet of the plasma membrane, the Stargardt disease-associated ABCA4 protein is a flippase responsible for importing retinoids into rod photoreceptors and retinal pigment epithelial (RPE) cells (Lenis et al., 2018; Sun et al., 1999; Weng et al., 1999). Thus, mutations in *ABCA4* result in accumulation of toxic retinoids and subsequent photoreceptor degeneration. In *Drosophila*, two ABCA proteins, Engulfment ABC Transporter in the ovary (Eato) and Lipid droplet defective (Ldd), export toxic lipids induced by oxidative stress from photoreceptors to nearby glia (Liu et al., 2015; Moulton et al., 2021). Consequently, the loss of *Eato* or *ldd* results in early photoreceptor degeneration in the presence of oxidative stress (Moulton et al., 2021). Despite these advances, whether ABCA proteins are involved in neurodegeneration through other means remains to be explored.

Besides their structural roles in membranes, lipids can also contribute to neurodegeneration through signaling functions. Phosphatidylserine (PS) is a phospholipid normally found in the inner leaflet of the plasma membrane (Segawa and Nagata, 2015). However, in sick or degenerating neurons, PS translocates to the extracellular leaflet, where it functions as a cell-surface “eat-me” signal to induce phagocytosis of neurites by nearby phagocytes (Sapar and Han, 2019). PS-induced phagocytosis not only enables clearance of neuronal debris resulting from degeneration but can also potently break down neurites of live neurons (Ji et al., 2022; Sapar et al., 2018). The asymmetric distribution of PS on the plasma membrane of healthy cells is established and maintained by flippases that belong to the P4-ATPase family of lipid transporters (Segawa et al., 2014; Segawa and Nagata, 2015). On the other hand, lipid scramblases in the TMEM16 and XKR families disrupt PS asymmetry and are responsible for PS exposure on platelets and apoptotic cells, respectively (Fujii et al., 2015; Suzuki et al., 2013; Suzuki et al., 2010).

Among ABCA proteins, murine ABCA1 was first found to promote Ca^2+^-induced PS exposure at the plasma membrane (Hamon et al., 2000). Similarly, a *C. elegans* ABCA protein, CED-7, was later discovered to facilitate PS exposure on apoptotic cells in the developing embryo and the subsequent transfer of PS-exposing extracellular vesicles from apoptotic cells to phagocytes (Mapes et al., 2012; Venegas and Zhou, 2007; Zhang et al., 2012). These observations are consistent with most ABCA proteins being lipid floppases (Phillips, 2018; Quazi and Molday, 2011), and their ability to promote PS exposure has been thought to be important for the functions of ABCA proteins in phagocytosis (Hamon et al., 2000). To date, it remains unknown how ABCA proteins may participate in neurodegeneration by regulating PS transport.

In this study, we show that the *Drosophila* ABCA protein Eato regulates phagocytosis-driven dendrite and axon degeneration by playing opposing roles in neurons and phagocytes: In neurons, it prevents dendrites and axons from being engulfed by phagocytes; in phagocytes, instead of being required for phagocytosis, it makes the cell more sensitive to PS presented by nearby engulfment targets. Thus, although the loss of *Eato* in neurons alone results in degeneration of diverse neurons in both the peripheral and central nervous systems, removing *Eato* in both neurons and phagocytes rescues neuronal degeneration. Despite these two distinct cell type-specific roles for Eato, unexpectedly, it functions in both neurons and phagocytes by suppressing, rather than enhancing, the effects of PS on cell surface. In support of Eato’s function in phagocytes, we further found that PS exposure on phagocytes inhibits, instead of promoting, phagocytosis. Finally, we show that CED-7 and several mammalian ABCA homologs can partially compensate for the loss of *Eato* in phagocytes, but not in neurons, suggesting both conserved and cell type-specific functions of ABCA proteins in regulating lipid homeostasis and phagocytosis-dependent neurodegeneration.

## RESULTS

### *Eato* LOF in da neurons causes engulfment-dependent dendrite degeneration

To search for lipid transporters that may affect exposure of eat-me signals on the plasma membrane of neurons, we screened candidate genes in the ABCA subfamily by RNA interference (RNAi) and CRISPR-induced mutagenesis (Poe et al., 2019) in *Drosophila* class IV dendritic arborization (C4da) neurons. C4da neurons are somatosensory neurons that grow highly elaborated dendrites underneath epidermal cells on the larval body wall (Grueber et al., 2002); these neurons are a well-established model system for studying degeneration and phagocytosis of dendrites (Han et al., 2014; Ji et al., 2022; Ji et al., 2023; Sapar et al., 2018). Among the genes we examined, only the loss-of-function (LOF) of *Eato* led to dendrite degeneration. In these assays, C4da neurons were labeled by MApHS, a dual fluorescent membrane marker that contains both a pH-sensitive pHluorin and an acid-resistant tdTomato (tdTom) (Han et al., 2014). Degenerating dendrites are engulfed by larval epidermal cells, resulting in tdTom-labeled neuronal debris inside phagosomes that are dispersed in epidermal cells (Han et al., 2014). To knock out *Eato* in C4da neurons, we combined the C4da-specific *ppk-Cas9* (Poe et al., 2019) with *gRNA-Eato*, which expresses ubiquitously two guide RNAs (gRNAs) targeting the shared coding sequence of both *Eato* splicing variants (Figure S1). Compared to control neurons that showed no debris (Figures 1A and 1F), *Eato* knockout (KO) neurons exhibited a 70.8% reduction in dendrite length along with widespread dendrite debris in the epidermis at 96 h after egg laying (AEL), indicating severe degeneration (Figures 1B, 1E and 1F). This phenotype was further confirmed by C4da-specific *Eato* knockdown (KD) with a short hairpin RNA (shRNA) transgene (HMC06027) targeting the shared coding sequence of *Eato* (Figures 1C–1F and S1). To understand how this degeneration phenotype develops, we examined *Eato* KO neurons at multiple developmental stages from 48 h AEL to 120 h AEL. *Eato* KO neurons did not show obvious signs of degeneration at 48 h AEL but displayed significant debris and gradually more severe dendrite reduction from 72 h AEL onward (Figures 1G–L), demonstrating a progressive loss of dendrites due to degeneration.

**Figure 1.**
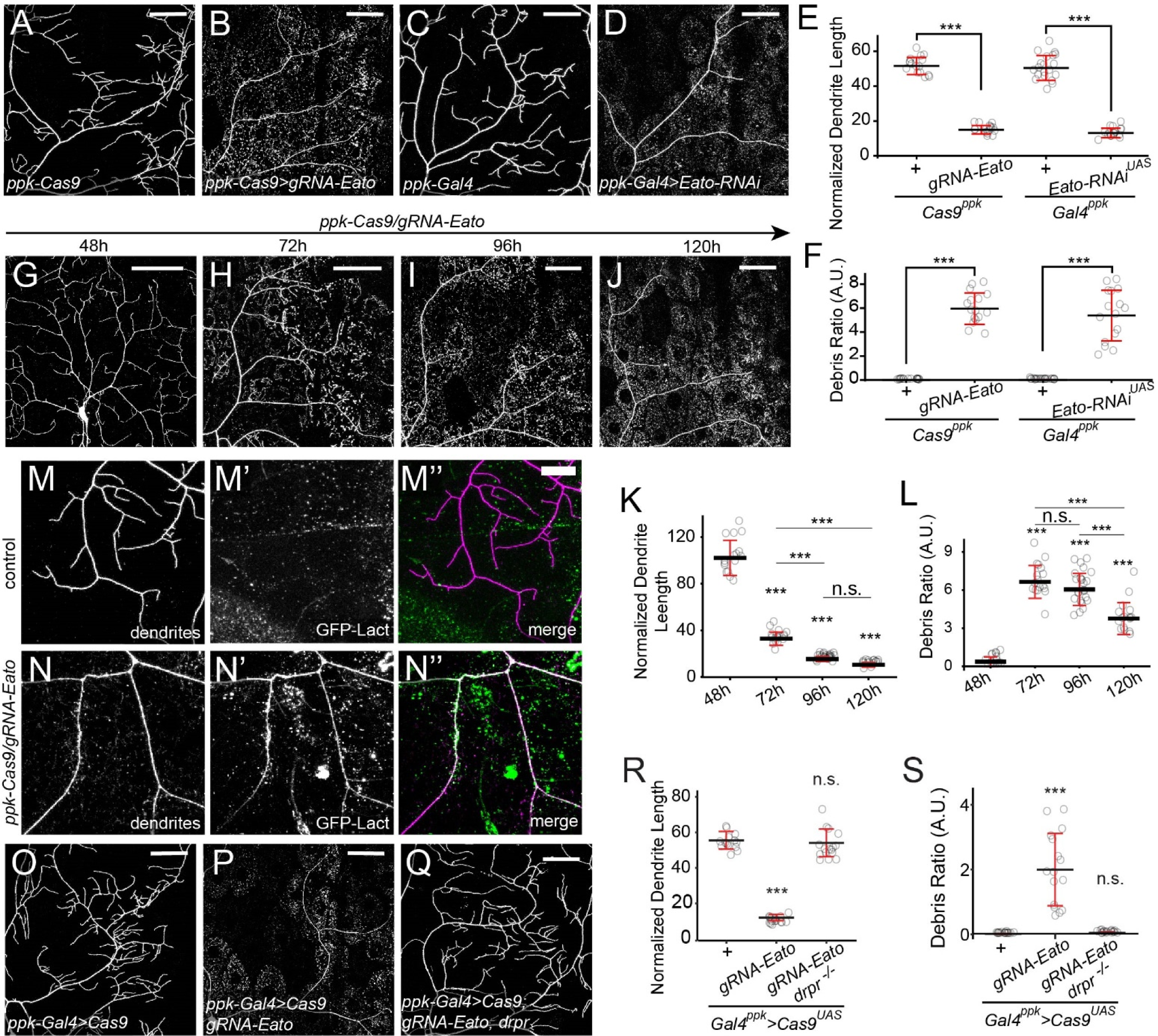
*Eato* LOF in C4da neurons leads to engulfment-dependent dendrite degeneration. (A–B) Dendrites of C4da neurons in *ppk-Cas9* control (A), C4da-specific *Eato* knockout (KO) (B), *ppk-Gal4* control (C), and C4da-specific *Eato* knockdown (KD) late 3^rd^ instar larvae. (E–F) Quantification of normalized dendrite length (total dendrite length/total area) (E) and debris ratio (debris area/total area) (F) in A–D. n = number of neurons and N = number of animals: *Cas9^ppk^* (n = 16, N = 9); *Cas9^ppk^ gRNA-Eato* (n = 16, N = 8); *Gal4^ppk^* (n = 20, N = 11); *Gal4^ppk^ Eato-RNAi^UAS^* (n = 16, N = 14). (G–J) Dendrites of C4da-specific *Eato* KO across different developmental stages. (K–L) quantification of normalized dendrite length (K) and debris ratio (L) in G–J. n = number of neurons and N = number of animals: 48h (n = 16, N = 8); 72h (n = 17, N = 11); 96h (n = 24, N = 12); 120h (n = 16, N = 8). (M–N’’) Binding patterns of the PS sensor GFP-Lact on dendrites of control (M–M’’) and *Eato* KO neurons (N–N’’) in late 3^rd^ instar larvae. Fat body-specific *dcg-Gal4* drives expression of GFP-Lact. (O–Q) C4da neuron dendrites in *ppk-Gal4 UAS-Cas9* control (O), C4da-specific *Eato* KO (P), and C4da-specific *Eato* KO in *drpr^indel3^* homozygous mutant late 3^rd^ instar larvae. (R–S) quantification of normalized dendrite length (R) and debris ratio (S) in O–Q. n = number of neurons and N = number of animals: *Gal4^ppk^* > *Cas9^ppk^* (n = 15, N = 10); *Gal4^ppk^* > *Cas9^ppk^ gRNA-Eato* (n = 16, N = 11); *Gal4^ppk^* > *Cas9^ppk^ gRNA-Eato drpr^indel3^* (n = 16, N = 12). C4da neurons were labeled by *ppk-MApHS* in (A–D) and (G–J), *ppk-CD4-tdTomato* in (M–N”), and *ppk-Gal4 UAS-CD4-tdTomato* in (O–Q). Scale bars: 50 μm in (A–D), (G–J), and (O–Q); 20 μm in (M– N”). In all plots, \*\*\**p*<0.001; n.s., not significant; one-way ANOVA with Tukey post-hoc test.

Degenerating neurons display the eat-me signal PS on their surface, which triggers phagocytic clearance of neuronal debris by phagocytes (Sapar et al., 2018). Considering that *Eato* encodes a putative ABCA lipid transporter, we wondered if *Eato* KO neurons display PS exposure. We previously developed an *in vivo* extracellular PS labeling system in which PS binding probes fused to fluorescent proteins, such as GFP-Lact, are expressed by the fat body and secreted into the larval body fluid (Sapar et al., 2018). Peripheral tissues with surface PS exposure are coated by the probes. Using GFP-Lact, we detected strong PS externalization on the dendrites of *Eato* KO neurons but not on control neurons (Figures 1M–1N’’), consistent with the degenerating state of these KO neurons.

We previously found that PS exposure on neuronal surfaces can induce neurite degeneration (Ji et al., 2022; Sapar et al., 2018). In such a scenario, phagocytes engulf PS-exposing (but intact and living) neurites, and the phagocytosis is responsible for the neurite degeneration. However, the observation of PS exposure on *Eato* KO neurons does not necessarily indicate that phagocytosis causes the degeneration, considering that PS exposure could be a consequence of membrane disruptions expected of dendrite degeneration (Sapar et al., 2018). To determine if the dendrite degeneration associated with *Eato* LOF depends on phagocytosis, we knocked out *Eato* in a null mutant of *draper* (*drpr*), which encodes an engulfment receptor required for larval epidermal cells to phagocytose degenerating dendrites (Han et al., 2014). Strikingly, dendrite degeneration of *Eato* KO neurons was completely suppressed in the *drpr* mutant (Figures 1P–1S). These data show that dendrite degeneration of *Eato* deficient neurons is caused by the phagocytic activity of epidermal cells.

### *Eato* LOF makes epidermal cells insensitive to degenerating dendrites

To further investigate the LOF phenotype of *Eato*, we generated an *Eato* CRISPR mutant by knocking out *Eato* in the germline. *Eato^10^* contains a deletion of 709 nucleotides from exon 4 to exon 6 between the two gRNA target sites (Figure S1), resulting in a reading-frame shift from amino acid 201, and thus is expected to be a null allele. Surprisingly, when examining C4da neurons in heterozygotes of *Eato^10^* and *Df(BSC812)*, a deficiency lacking the entire *Eato* locus, we did not observe any signs of dendrite degeneration (Figures 2B, 2E and 2F), a sharp contrast with the C4da-specific KO of *Eato* (Figures 2A, 2E and 2F). *Eato* was previously shown to be involved in phagocytosis of nursing cells by follicular epithelial cells during *Drosophila* oogenesis (Santoso et al., 2018), and ABCA homologs in worms (CED-7) and mammals (ABCA1) participate in phagocytosis as well (Hamon et al., 2000; Venegas and Zhou, 2007). Given the requirement of phagocytosis for the dendrite degeneration of *Eato*-deficient neurons (Figures 1P–1S), we hypothesized that epidermal cells require *Eato* to engulf *Eato*-deficient dendrites. To test this idea, we knocked down *Eato* simultaneously in both neurons and epidermal cells. Indeed, this led to a wildtype (WT) dendrite phenotype (Figures 2D–2F), as compared to the severe degeneration seen in neuron-specific KD (Figures 2C, 2E and 2F), confirming a requirement for *Eato* in the phagocytic destruction of *Eato* mutant neurons by epidermal cells.

**Figure 2.**
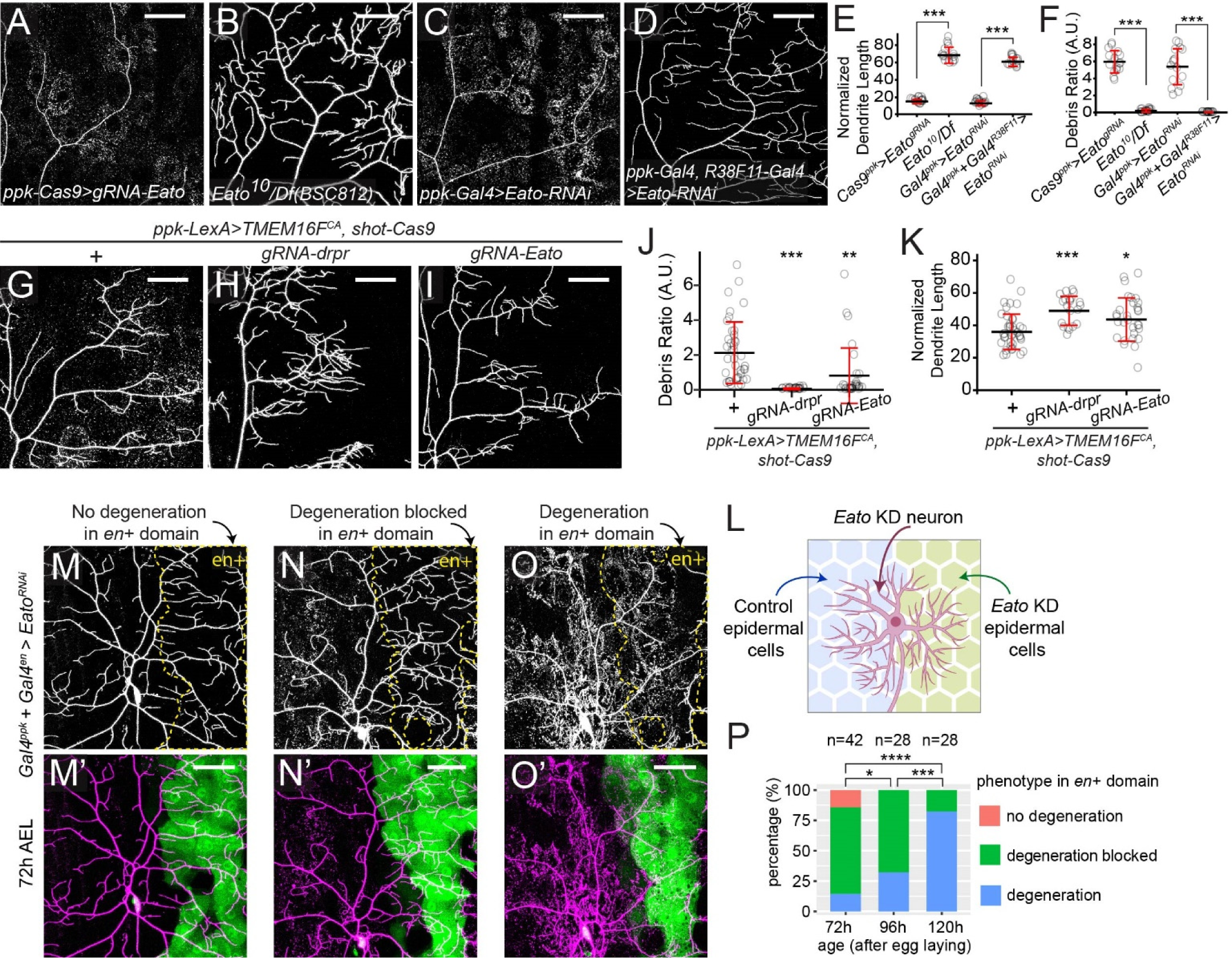
*Eato* LOF makes epidermal cells insensitive to degenerating dendrites. (A–D) Dendrites of C4da neurons in C4da-specific *Eato* KO (A), *Eato^10^/Df(BSC812)* transheterozygous mutant (B), C4da-specific *Eato* KD (C), and both C4da and epidermal cell-specific *Eato* KD (D) late 3^rd^ instar larvae. (E–F) Quantification of normalized dendrite length (E) and debris ratio (F) in (A–D). n = number of neurons and N = number of animals: *Cas9^ppk^* > *Eato^gRNA^* (n = 16, N = 8); *Eato^10^/Df* (n = 16, N = 7); *Gal4^ppk^* > *Eato^RNAi^* (n = 16, N = 14); *Gal4^ppk^ + Gal4^R38F11^ > Eato^RNAi^* (n = 16, N = 9). (G–I) C4da neuron dendrites of TMEM16F^CA^-overexpressing (OE) C4da neurons with control epidermal cells (G), epidermal cell-specific *drpr* KO (H), and epidermal cell-specific *Eato* KO (I) in late 3^rd^ instar larvae. (J–K) Quantification of normalized dendrite length (J) and debris ratio (K) in (G–I). n = number of neurons and N = number of animals: *ppk-LexA>TMEM16F^CA^ shot-Cas9* (n = 40, N = 19); *ppk-LexA>TMEM16F^CA^ shot-Cas9 gRNA-drpr* (n = 21, N = 15); *ppk-LexA>TMEM16F^CA^ shot-Cas9 gRNA-Eato* (n = 30, N = 15). (L) A diagram of experiment design in (M–O’). *Eato* is knocked down in C4da neurons (by *ppk-Gal4* driver) and the posterior epidermal cells of each segment (by *en-Gal4* driver). The anterior epidermal cells where Gal4 is not expressed serve as an internal control. (M–O’) Dendrites of C4da neurons in 72 h AEL animals from the experiment described in (L). Binary images (M, N, O) show dendrites only and color images show neurons and *en+* domain (M’, N’, O’). *en+* domains in binary images are enclosed by yellow dashed lines. Three categories of phenotypes were observed: no degeneration (M–M’), degeneration blocked in the *en+* domain (N–N’), and degeneration in *en+* domain (O-O’). (P) Quantification of percentage of three indicated genotypes in 72 h to 120 h AEL animals. n = number of neurons and N = number of animals: 72h (n = 42, N = 18); 96h (n = 28, N = 14); 120h (n = 28, N = 14). Pan-epidermal expression is driven *R38F11-Gal4*. Pan-epidermal KO is driven by *shot-Cas9*. C4da neurons were labeled by *ppk-MApHS* in (A–D), (G–I) and (M–O’). Scale bars: 50 μm. In all plots, *p<0.05, ***p<0.001, ****p<0.0001; n.s., not significant. (E–F) and (J–K), one-way ANOVA with Tukey post-hoc test; (P), chi-square test, *p*-value adjusted by FDR method.

*Eato* could be required for epidermal cells to engulf *Eato*-deficient neurons specifically or to engulf any neuron displaying eat-me signals. To distinguish between these two possibilities, we sought to induce ectopic PS exposure on dendrites using the murine TMEM16F, a calcium-dependent phospholipid scramblase whose activity results in externalization of PS on the plasma membrane (Suzuki et al., 2010). Because expression of TMEM16F in C4da neurons causes only weak PS exposure (Sapar et al., 2018), to induce stronger PS exposure on dendrites, we expressed a constitutively active TMEM16F mutant (TMEM16F^CA^) carrying two mutations (Y563K/D703R) that make the scramblase calcium-independent (Le et al., 2019). As expected, TMEM16F^CA^ expression resulted in a moderate level of dendrite debris in epidermal cells (Figures 2G and 2J). The dendrite debris was completely eliminated, however, when *drpr* was simultaneously knocked out from epidermal cells using tissue-specific CRISPR (Poe et al., 2019) (Figures 2H and 2J), confirming the dependence of this dendrite degeneration on phagocytosis. Importantly, epidermal-specific *Eato* KO also suppressed the dendrite debris of TMEM16F^CA^-expressing neurons and increased the dendrite length (Figures 2I–2K), even though in either case not as potently as *drpr* KO. These results suggest that *Eato* LOF in epidermal cells impairs the ability of these cells to engulf PS-exposing dendrites.

To distinguish whether *Eato* is strictly required for phagocytosis or simply enhances the ability of epidermal cells to engulf degenerating neurons, we knocked down *Eato* in epidermal cells and severed C4da dendrites using laser. We previously showed that physical severing of dendrites induces rapid and high PS exposure on the detached dendrites within a few hours, which drives engulfment and fragmentation of the dendrites (Ji et al., 2022; Sapar et al., 2018) (Figures S2A and S2D). The engulfment of such injured dendrites is blocked in *drpr* mutants (Han et al., 2014; Ji et al., 2022), as reflected by dendrite fragments lined up in the original dendritic pattern 20 h after injury (AI) (Figure S2B). In contrast, injured dendrites were completely engulfed in *Eato* KD, resulting in widely spread dendrite debris in each epidermal cell (Figures S2C–S2E). These data suggest that *Eato* is not required for phagocytosis per se; rather, it is needed for epidermal cells to detect moderate levels of PS exposure on dendrites, such as in TMEM16F^CA^ expression.

To further confirm that Eato only enhances phagocyte sensitivity but is not required for engulfment, we exposed *Eato*-deficient neurons (via KD) to both WT and *Eato*-deficient epidermal cells in the same animal. The WT control epidermal cells are in the anterior half of each segment, while *Eato* KD is induced in the posterior half of the segment by *en*-*Gal4* (Figure 2L). We reasoned that the anterior WT epidermal cells should attack *Eato*-deficient neurons and cause dendrite injury. If the injury signals spread to posterior dendrites, these dendrites may show elevated levels of eat-me signals and thus allow us to test the ability of *Eato*-deficient posterior epidermal cells to engulf them. In 3^rd^ instar animals, we observed three classes of phenotypes: (1) a “no degeneration” phenotype, in which neither the control domain (anterior) nor the *en*+ domain (posterior) showed dendrite degeneration (Figures 2M and 2M’); (2) a “blocked degeneration” phenotype, in which only the control but not the *en*+ domain showed dendrite degeneration (Figures 2N and 2N’); and (3) a “degeneration” phenotype, in which both the control and the *en*+ domain showed dendrite degeneration (Figures 2O and 2O’). At 72 h AEL, most neurons (71.4%) showed “blocked degeneration”, and only small fractions showed “no degeneration” (14.3%) or “degeneration” (14.3%) (Figures 2P), suggesting that posterior dendrites are locally protected due to *Eato* LOF in posterior epidermal cells. However, the “degeneration” phenotype increased to 32.1% at 96 h AEL and 82.1% at 120 h AEL (Figures 2P). These observations are consistent with the idea that posterior dendrites become increasingly sick with age and are eventually recognized and engulfed by *Eato*-deficient epidermal cells. Thus, *Eato* is not necessary for phagocytosis in epidermal cells, but the loss of *Eato* reduces the sensitivity of epidermal cells to mildly unhealthy dendrites and delays the initiation of phagocytosis.

### *Eato* LOF in da neurons causes glia-dependent axon degeneration

Like dendrites, axons exhibit PS-induced engulfment and degeneration (Kim et al., 2010; Sapar et al., 2018; Shacham-Silverberg et al., 2018). However, axons and dendrites differ in their requirements for specific components of the axon-death pathway in injury-induced degeneration (Ji et al., 2022). We thus tested if *Eato* is also required to maintain the integrity of C4da axons, which project to the ventral nerve cord (VNC) in a ladder pattern (Figure 3A). When *Eato* was knocked out from C4da neurons, the axon ladder became fragmented (Figures 3B and 3D), indicating axon degeneration. Like dendrite degeneration, this axon degeneration was also completely suppressed in the *drpr* mutant (Figures 3C and 3D), demonstrating that phagocytosis also drives axon degeneration of *Eato*-deficient neurons.

**Figure 3.**
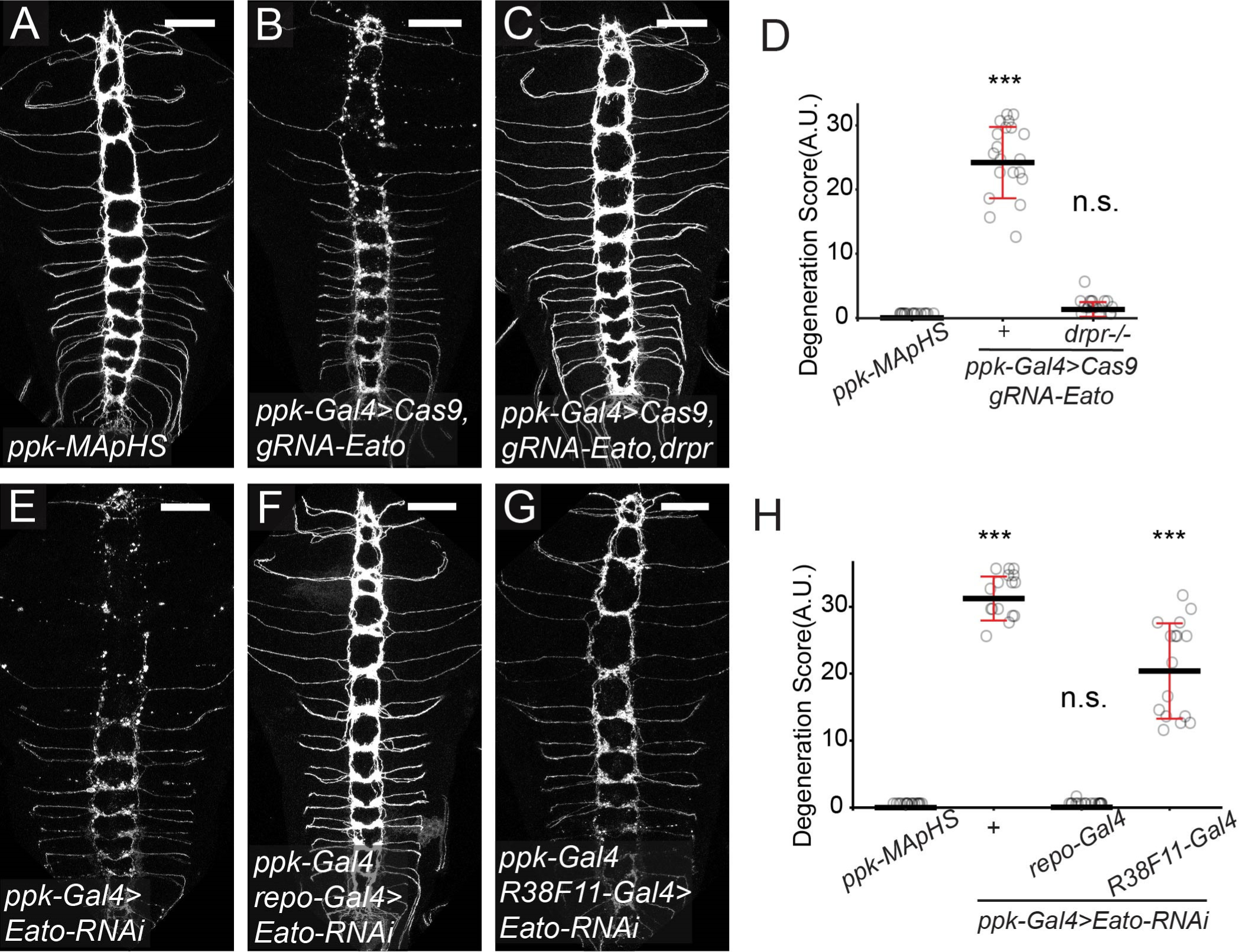
*Eato* LOF in da neurons causes glia-dependent axon degeneration. (A–D) Axons of C4da neurons in *ppk-Gal4 UAS-Cas9* control (A), C4da-specific *Eato* KO (B), and C4da-specific *Eato* KO in *drpr^indel3^* homozygous mutant (C) late 3rd instar larvae. Degeneration level is quantified as degeneration score in (D) (see Materials and Methods). n = number of brains: *ppk-MApHS* (n = 16); *ppk-Gal4>Cas9 gRNA-Eato* (n = 21); *ppk-Gal4>Cas9 gRNA-Eato drpr^-/-^* (n = 19). (E–H) Axons of C4da neurons in C4da-specific *Eato* KD (E), C4da and glia-specific *Eato* KD (F), and C4da and epidermal cell-specific *Eato* KD (G) animals. Degeneration level is quantified as degeneration score in (H). n = number of brains: *ppk-MApHS* (n = 16); *ppk-Gal4>Eato-RNAi* (n = 17); *ppk-Gal4 + repo-Gal4 >Eato-RNAi* (n = 18), *ppk-Gal4 + R38F11-Gal4 >Eato-RNAi* (n = 16). Glia-specific expression is driven by *repo-Gal4*. C4da neurons were labeled by *ppk-MApHS* in (A–C) and (E–G). Scale bars: 50 μm. In all plots, \*\*\**p*<0.001; n.s., not significant, one-way ANOVA with Tukey post-hoc test.

In the central nervous system (CNS), glia are the phagocytes that engulf dead neurons and neuronal debris (Freeman, 2015; Sapar and Han, 2019). The axons of da neurons are surrounded by glia in both peripheral nerves and the VNC. Hence, we asked whether glia are involved in axon degeneration of *Eato*-deficient neurons by knocking down *Eato* in both C4da neurons and glia. Unlike C4da-specific *Eato* KD, which showed drastic axon degeneration (Figures 3E and 3H), *Eato* KD in both neurons and glia showed no axon degeneration (Figures 3F and 3H). These data confirm that glia are required for the phagocytosis-dependent axon degeneration and further demonstrate that *Eato* promotes phagocytic activity of glia.

Although the dendritic and axonal compartments of the same da neuron are attacked by different phagocyte types, we wondered if degeneration of the two compartments is coupled. We thus knocked down *Eato* in both neurons and epidermal cells to suppress dendrite degeneration (Figures S3A, S3C, S3E and S3F) and examined axon morphology. Interestingly, these neurons showed much weaker axon degeneration than neuronal KD alone (Figures 3E, 3G and 3H). In contrast, knocking down *Eato* in both neurons and glia to suppress axon degeneration had no impact on dendrite degeneration (Figures S3D and S3F). These data together suggest that damage to dendrites strongly enhances phagocytosis of axons, likely by promoting exposure of eat-me signals on axons, but axon degeneration is compartmentally restricted and does not affect dendrites in the same neuron.

### *Eato* encodes a membrane protein required for the integrity of diverse neurons in both the peripheral and central nervous systems

To determine where else *Eato* may play a role in protecting neurons from degeneration, we first examined Eato expression patterns by generating an *Eato-Gal4* transcription reporter. *MiMIC^MI14571^* (Venken et al., 2011) is a transgenic insertion in the second intron, which is between two coding exons shared by both *Eato* isoforms (Figure S1). We converted *MiMIC^MI14571^* to a 2A-Gal4 Trojan exon through recombinase-mediated cassette exchange (Figure S4A) (Diao et al., 2015). The resultant *Eato-Gal4* should produce a short truncated Eato protein (102 amino acids) and, more importantly, a Gal4 driven by the endogenous regulatory sequence of *Eato* that should recapitulate the *Eato* expression pattern.

By crossing to UAS-driven fluorescent reporters, we observed broad *Eato-Gal4* expression in peripheral tissues, including da neurons, bipolar dendrite (bd) neurons, a subset of external sensory (es) neurons, epidermal cells (Figures 4A–4A”), peripheral glia (Figures 4B–4B”), muscles (Figures S4B and S4C), and the trachea (Figure S4D). In the VNC, Eato-Gal4 expression overlaps with some but not all neurons and glia (Figures 4C–4D”).

**Figure 4.**
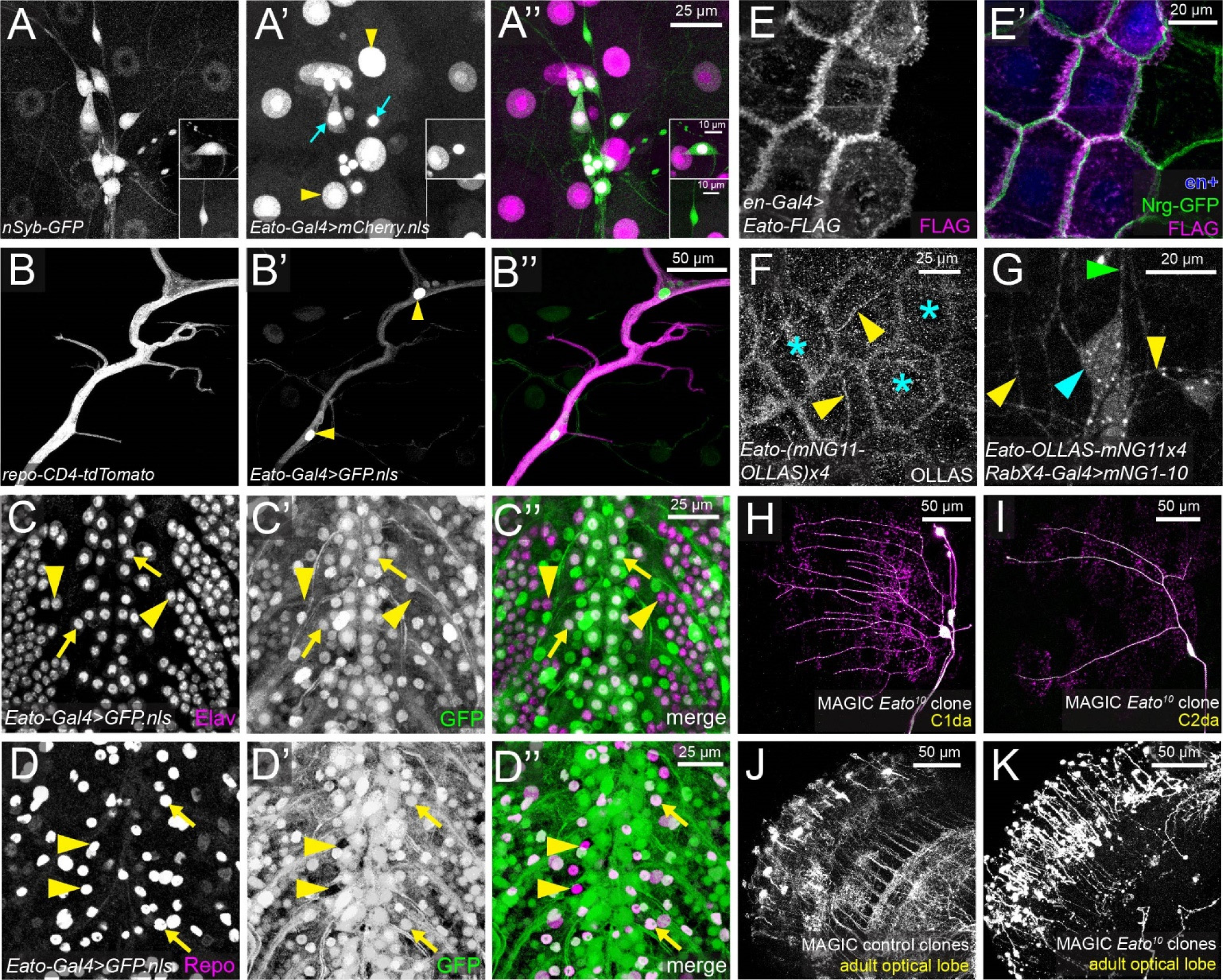
*Eato* encodes a membrane protein required for the integrity of diverse neurons in both PNS and CNS. (A–A’’) *Eato* expression pattern on the body wall in a 96 h AEL larva. Neurons are labeled by *nSyb-tdGFP* (A). *Eato-Gal4^MI14571^* drives the expression of a nuclear mCherry (A’). Yellow arrowheads: epidermal nuclei; blue arrows: neuronal nuclei. Upper insets show an es neuron expressing *Eato* and lower insets show an es neuron without *Eato* expression. (B–B’’) *Eato* expression in peripheral glial cells, showing an intersegmental nerve bundle. Glial cells are labeled by *repo-CD4-tdTomato* (B). *Eato-Gal4^MI14571^* drives the expression of a nuclear GFP (B’). Yellow arrowheads: glial nuclei. (C–D’’) *Eato* expression pattern in the central nervous system (CNS). *Eato-Gal4^MI14571^* drives the expression of a nuclear GFP (C’ and D’). (C–C’’) show Elav staining to visualize neuronal nuclei. Arrows: neurons with *Eato* expression; arrowheads: neurons without *Eato* expression. (D–D’’) show Repo staining to visualize glial nuclei. Arrows: glia with *Eato* expression; arrowheads: glia without *Eato* expression. (E–E’) Localization of FLAG-tagged Eato protein in epidermal cells. Eato(B) is expressed by *en-Gal4* and detected by FLAG antibody staining. In (E’), the *en*+ domain is visualized by mIFP expression (blue), and cell junctions are indicated by Nrg-GFP (green). (F) Endogenous Eato in a homozygous *Eato-(mNG_11_-OLLAS)_X4_* knock-in larva as detected by OLLAS staining. Yellow arrowheads: dendrite tracks; blue asterisks: epidermal cells. (G) Endogenous Eato protein in neurons of homozygous *Eato-(mNG_11_-OLLAS)_X4_* detected by split-mNeonGreen reconstitution. mNG_1-10_ is expressed in all neurons by *RabX4-Gal4*. Cyan arrowhead: soma; green arrowhead: axon; yellow arrowheads: dendrites. (H–I) *Eato^10^* homozygous clones in class I da (C1da) (H) and class II da (C2da) (I) neurons generated by the MAGIC method. Neuronal clones were labeled by *RabX4-Gal4 UAS-MApHS*. pHluorin is in green and tdTom is in magenta. Magenta-only signals indicate neuronal debris engulfed by epidermal cells. (J–K) Control (J) and *Eato^10^* mutant (K) neuronal clones generated by the MAGIC method in the adult optical lobe. Neuronal clones were labeled by *RabX4-Gal4 UAS-MApHS* but only tdTom signal is shown.

Next, we asked where the Eato protein is localized in cells. First, we generated a FLAG-tagged *UAS-Eato* (the long isoform B) transgene and expressed it in epidermal cells. FLAG staining was detected strongly on the lateral plasma membrane, overlapping with the membrane marker Nrg-GFP, and also at lower levels in intracellular vesicles (Figures 4E and 4E’). Next, to determine the localization of endogenous Eato proteins in epidermal cells and neurons, we inserted a (mNG_11_-OLLAS)x4-2A-QF2 cassette in the *Eato* locus immediately before the stop codon of the longer isoform (Fig. S4E) using a gRNA-donor vector optimized for CRISPR activity in the *Drosophila* germline (Koreman et al., 2021). mNeonGreen_11_ (mNG_11_) is a fragment of split mNG and can reconstitute the full fluorescent mNG protein when mNG_1-10_ is co-expressed in the same cell (Feng et al., 2017). OLLAS is a short tag, for which high-affinity antibodies are available (Park et al., 2008). 2A-QF2 in the construct allows identification of knock-in (KI) candidates by crossing to *QUAS-*driven fluorescent reporters (Riabinina et al., 2015). mNG_11_ enables detection of Eato proteins in specific cells, while OLLAS staining can visualize Eato proteins in all expressing tissues. Using OLLAS staining, we confirmed the presence of Eato on the plasma membrane and intracellular vesicles of epidermal cells (stars in Figure 4F) and detected signals that appeared as dendritic patterns of sensory neurons (arrowheads in Fig. 4F). By expressing mNG_1-10_ using a pan-neuronal Gal4 (*RabX4-Gal4*), we detected reconstituted mNG signals on the soma, axons, and dendrites of da neurons (Figure 4G). mNG fluorescence appeared as smooth signals on the neuronal surface and also in bright puncta resembling intracellular vesicles.

The broad expression of *Eato* in the nervous system prompts the question of whether *Eato* is also important in neurons other than C4da. To answer this question, we generated MApHS-labeled *Eato^10^* homozygous mutant clones in both the peripheral nervous system (PNS) and the CNS of otherwise *Eato^10/+^* heterozygous animals using a technique called mosaic analysis by gRNA-induced crossing-over (MAGIC) (Allen et al., 2021). *Eato^10^* clones of class I-III da neurons showed severe dendritic degeneration as indicated by reduced dendrites and extensive neuronal debris near dendrites (Figures 4H, 4I and S4F). Mutant multi-dendritic dmd1 neurons also showed debris near dendrites (Figure S4G). Interestingly, we did not observe obvious degeneration at axon terminals of motor neurons (Fig. S4H). In the larval VNC (Figures S4I and S4J), the larval brain (Figures S4K and S4L), adult optical lobe (Figures 4J and 4K), and central brain (Figures S4M and S4N), *Eato^10^* mutant neurons showed severely fragmented neurites with blebbing, contrasting with the smooth and continuous signals on WT neuronal clones. These data show that *Eato* is required to maintain the integrity of a broad range of neurons. Consistent with this conclusion, pan-neuronal *Eato* KD using *RabX4-Gal4* caused pupal lethality. Considering that homozygous *Eato* mutant strains are viable and fertile, degeneration of *Eato* homozygous mutant neurons in otherwise heterozygous animals is most likely caused by *Eato*-dependent engulfment activity of resident phagocytes in the PNS and CNS, as is the case for C4da neurons (Figure 2D).

### Putative ABCA transporter activity is required for Eato’s function

The *Eato* locus produces two alternatively spliced transcripts of different lengths (Fig. S1): The longer *Eato-RB* isoform encodes a full-length ABCA protein, with two ABC transporter-like ATP-binding domains (InterPro IPR003439), while the shorter *Eato-RC* isoform ends before the first ABC transporter-like domain. To determine if the full-length isoform is responsible for Eato’s function in neurons and phagocytes, we knocked down *Eato* using two additional RNAi lines (GD1133 and KK104197) that target *Eato-RB* only (Figure S1). Expression of *Eato-RNAi^KK104197^* in C4da neurons with *ppk-Gal4* caused strong dendrite degeneration, albeit slightly weaker than that with *Eato-RNAi^HMC06027^* (Figures 5A–5E). When driven by *Gal4^21-7^*, an early pan-da driver (Song et al., 2007), *Eato-RNAi^GD1133^* also caused strong dendrite degeneration in C4da neurons (Figures S5A–S5D). These data indicate that the *Eato-RB* isoform is necessary for neuronal maintenance.

**Figure. 5.**
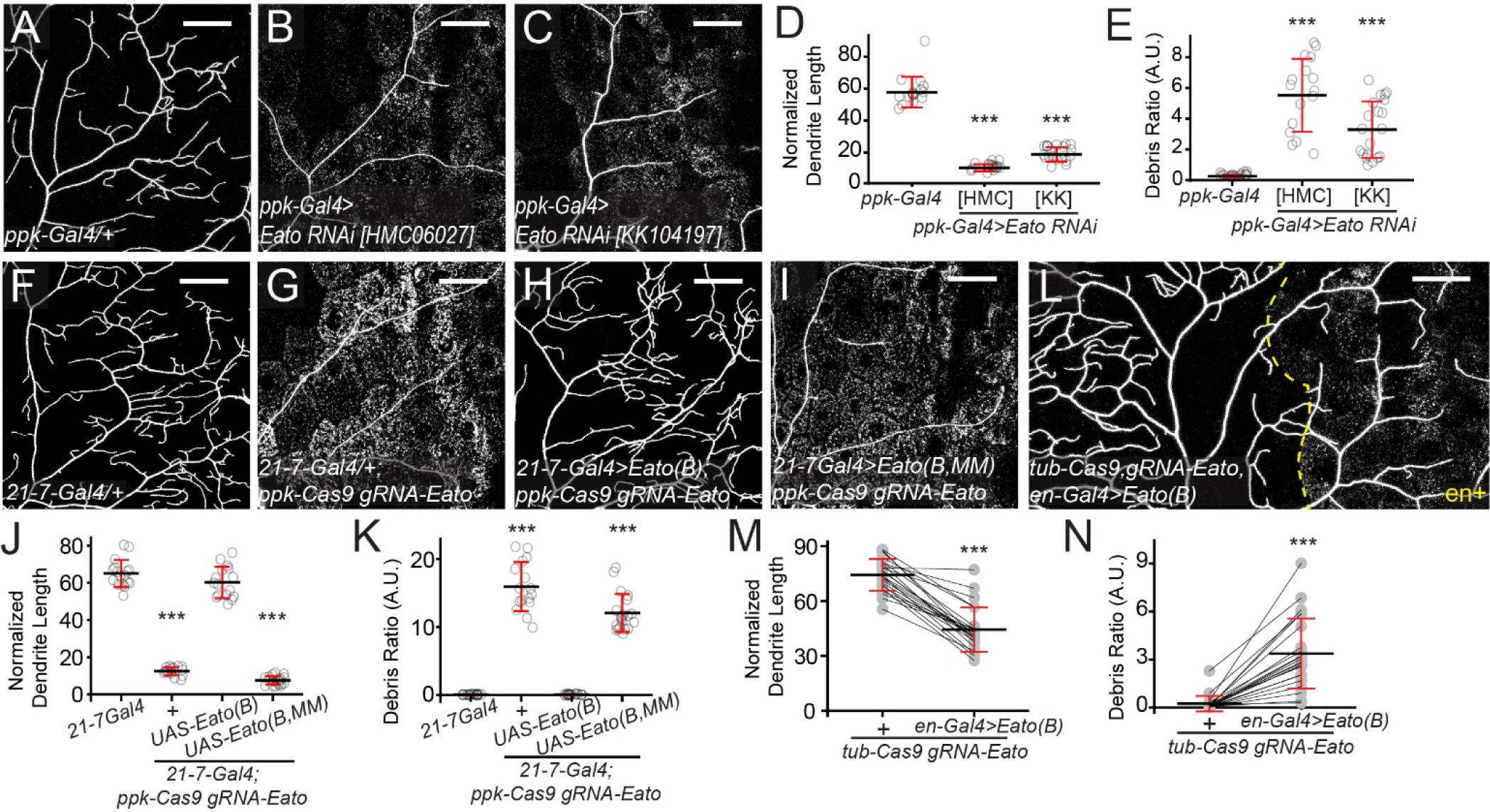
Putative ABCA transporter activity is required for Eato’s function. (A–C) Dendrites of C4da neurons in *ppk-Gal4* control (A), C4da-specific *Eato(B/C)* KD (B), and C4da-specific *Eato(B)* KD (C) late 3^rd^ instar larvae. (D–E) Quantification of normalized dendrite length (D) and debris ratio (E) of neurons in (A–C). n = number of neurons and N = number of animals: *ppk-Gal4* (n = 16, N = 11); *ppk-Gal4>Eato-RNAi[HMC]* (n = 16, N = 14); *ppk-Gal4>Eato-RNAi[KK]* (n = 20, N = 13). (F–K) Dendrites of C4da neurons in *21-7-Gal4* control (F), C4da-specific *Eato* KO (G), C4da-specific *Eato* KO with da-specific *Eato(B)* OE (H), and C4da-specific *Eato* KO with da-specific *Eato(B.MM)* OE (I) late 3^rd^ instar larvae. Normalized dendrite length is quantified in (J), and debris ratio is quantified in (K). n = number of neurons and N = number of animals: *21-7Gal4* (n = 16, N = 8); *21-7-Gal4 ppk-Cas9 gRNA-Eato* (n = 16, N = 8); *21-7-Gal4>UAS-Eato(B) ppk-Cas9 gRNA-Eato* (n = 17, N = 11), *21-7-Gal4>UAS-Eato(B,MM) ppk-Cas9 gRNA-Eato* (n = 20, N = 12). (L) Dendrites of C4da neurons in *en-Gal4 UAS-Eato(B); tub-Cas9 gRNA-Eato UAS-mIFP* animals. The *en+* domain is right to the yellow dashed line. The anterior non-expressing region serves as a control. Comparisons between the control (+) and *en*+ domains are shown in (M) (for normalized dendrite length) and (N) (for debris ratio). Data is from 23 neurons in 14 animals. *21-7-Gal4* drives expression in da neurons. C4da neurons were labeled by *ppk-MApHS* in (A–C), (F–I) and (L). Scale bars: 50 μm. In all plots, ***p<0.001; n.s., not significant. (D–E) and (J–K), one-way ANOVA with Tukey post-hoc test. (M–N), paired t-test.

Next, to determine if the long Eato protein isoform is sufficient for *Eato*’s function, we expressed *UAS-Eato(B)* in *Eato* KO animals. The gRNA target sequences in the *UAS-Eato(B)* coding sequence were altered by silent mutations to make the transgene gRNA-resistant. Eato(B) overexpression does not cause any dendrite phenotypes in WT neurons (Figures S5E–S5H), and it completely prevented dendrite degeneration caused by *Eato* KO (Figures 5F–5H, 5J and 5K), demonstrating the sufficiency of the long isoform in maintaining neuronal integrity. Furthermore, to determine if the putative ATPase activity of Eato is needed for its function, we mutated the key lysine (K) residue of the Walker A motif in each of the ATP-binding domains of Eato(B) into methionine (M). The resulting Eato(B.MM) mutant protein is predicted to be incapable of binding ATP and to lack transporter function (Anderson and Welsh, 1992; Hamon et al., 2000). In contrast to *UAS-Eato(B)*, *UAS-Eato(B.MM)* failed to rescue *Eato* KO neurons (Figures 5I–5K).

Lastly, to test if Eato(B) can restore the phagocytic activity of *Eato*-deficient epidermal cells, we expressed Eato(B) in epidermal cells of whole-body *Eato* KO animals. The KO was achieved by using a ubiquitously expressed *tub-Cas9*, and Eato(B) expression was driven by *en-Gal4*, so that non-expressing epidermal cells in the anterior hemi-segment can serve as an internal control. These animals displayed dendrite degeneration specifically in the posterior hemi-segment, as indicated by the reduced dendrite density and the presence of dendrite debris (Figures 5L–5N), suggesting successful rescue of epidermal phagocytic activity. As a comparison, overexpressing Eato(B) in WT epidermal cells did not enable them to engulf WT dendrites (Figures S5I–S5L).

Altogether, the above data indicate that the full ABCA sequence of Eato is necessary and sufficient for its function in neurons and phagocytes, and that the putative transporter activity is necessary for its function.

### Eato prevents dendrite degeneration by antagonizing neuronal PS exposure

Given the role of PS exposure in inducing phagocytosis and the PS exposure observed on *Eato* KO neurons, we wondered if the degeneration of *Eato*-deficient neurons is caused by PS exposure. To answer this question, we overexpressed *Drosophila* ATP8A in *Eato* KO neurons. ATP8A is a PS flippase that keeps PS in the inner leaflet of the plasma membrane (Ji et al., 2023) and its overexpression can suppress PS exposure in both neurons and phagocytes (Ji et al., 2022; Ji et al., 2023). Neuronal expression of ATP8A completely suppressed the degeneration of *Eato* KO neurons (Figures 6A–6D), suggesting that PS exposure is indeed the cause of the degeneration. This result also suggests that Eato’s normal function in neurons is to prevent PS exposure. We thus tested whether Eato(B) can antagonize ectopic PS exposure caused by disruptions of membrane lipid asymmetry.

**Figure 6.**
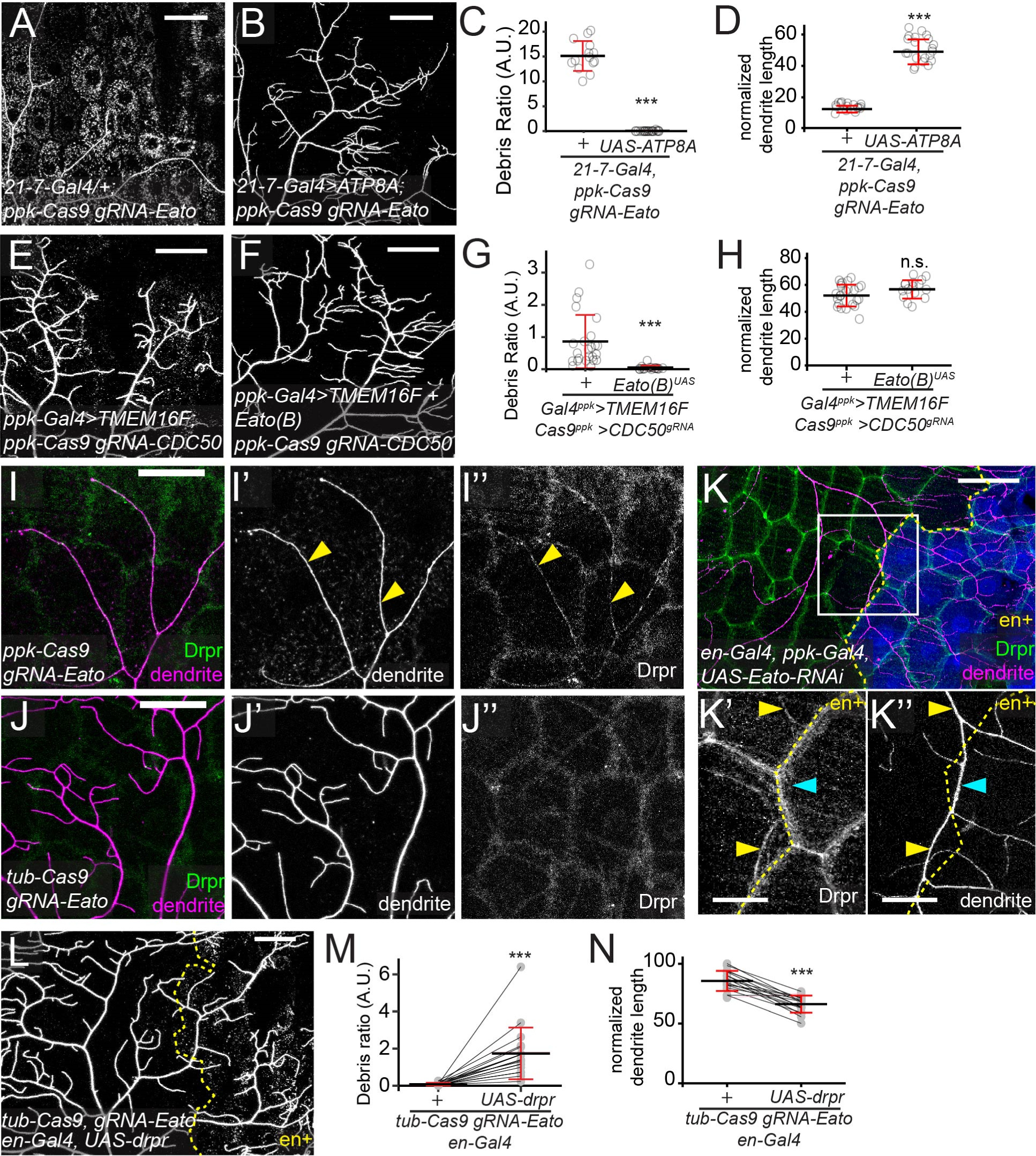
Eato suppresses PS exposure in neurons and facilitates Drpr recruitment to degenerating dendrites in epidermal cells. (A–D) Dendrites of C4da neurons in C4da-specific *Eato* KO (A) and C4da-specific *Eato* KO with da-specific ATP8A expression (B) late 3^rd^ instar larvae. Debris ratio is quantified in (C) and normalized dendrite length is quantified in (D). n = number of neurons and N = number of animals: *21-7-Gal4 ppk-Cas9 gRNA-Eato* (n = 16, N = 8); *21-7-Gal4 UAS-ATP8A ppk-Cas9 gRNA-Eato* (n = 21, N = 11). (E–H) Dendrites of C4da neurons with ectopic PS exposure (E) and additional Eato(B) overexpression (F) in late 3^rd^ instar larvae. Ectopic PS exposure was induced by TMEM16F overexpression and simultaneous *CDC50* KO. Debris ratio is quantified in (G) and normalized dendrite length is quantified in (H). n = number of neurons and N = number of animals: *Gal4^ppk^>TMEM16F Cas9^ppk^>CDC50^gRNA^* (n = 24, N = 14); *Gal4^ppk^>TMEM16F + Eato(B) Cas9^ppk^>CDC50^gRNA^* (n = 16, N = 12). (I–J’’) Drpr-GFP distribution in C4da-specific *Eato* KO (I–I’’) and whole-body *Eato* KO (J–J’’) mid-3^rd^ instar larvae. Yellow arrowheads: colocalization between dendrites and Drpr-GFP. (K–K’’) Drpr distribution in a mid-3^rd^ instar larva where *Eato* is knocked down in both C4da neurons and the *en*+ domain. The *en*+ domain is marked by mIFP (blue) in (K) and located right to the yellow dashed line in (K’ and K”). Drpr is detected by antibody staining. Yellow arrowheads: Drpr colocalized with dendrites in control epidermal cells; cyan arrowheads: absence of Drpr along dendrites in *Eato* KD epidermal cells. (L) Dendrites of C4da neurons in a whole-body *Eato* KO late 3^rd^ instar larva with Drpr overexpression by *en-Gal4*. The *en*+ region is right to the yellow dashed line. Debris ratio in control (+) and Drpr overexpressing epidermal cells is quantified in (M) and normalized dendrite length is quantified in (N). Data is from 18 neurons in 11 animals. C4da neurons were labeled by *ppk-MApHS* in (A–B) and (L); *ppk-CD4-tdTomato* in (E–F) and (I–K’’). Scale bars: 50 μm in (A–B), (E–F), (I–J), (K) and (L); 20 μm in (K’–K’’). In all plots, ***p<0.001; n.s., not significant. (C–D) and (G–H), two-sample t-test; (M–N), paired t-test.

Ectopic PS exposure in C4da neurons can be induced by overexpressing TMEM16F and simultaneous KO of *CDC50* (Sapar et al., 2018), which encodes an obligatory chaperone for ATP8A (Tanaka et al., 2011). Compared to these PS-exposing neurons, which were associated with moderate levels of neuronal debris (Figures 6E and 6G), additional Eato(B) expression in the neurons eliminated neuronal debris (Figures 6F and 6G). These results suggest that Eato normally suppresses PS exposure on neuronal surface to prevent neurites from being engulfed by phagocytes.

### Epidermal Eato facilitates Drpr recruitment to degenerating dendrites

The engulfment receptor Drpr is normally found only at low levels on the plasma membrane of phagocytes, but it is recruited to the site of engulfment in response to PS exposure on the engulfment target (Ji et al., 2023; MacDonald et al., 2006). To understand the engulfment defects of *Eato*-deficient epidermal cells, we examined Drpr recruitment to degenerating dendrites. As expected, distinct Drpr staining was detected along the degenerating dendrites of *Eato* KO neurons, in addition to the lateral membranes of epidermal cells (Figures 6I–6I”). However, the dendrite-overlapping Drpr staining was absent in whole-body *Eato* KO (Figures 6J–6J”). Considering that dendrites did not degenerate in whole-body *Eato* KO, the lack of Drpr accumulation on dendrites could be due to the absence of dendrite degeneration. To cause dendrites contacting *Eato*-deficient epidermal cells to degenerate, we knocked down *Eato* in both C4da neurons and *en*+ epidermal cells. In these animals, branches that straddle the border between WT and *Eato*-KD epidermal cells may undergo degeneration due to phagocytic attack by WT epidermal cells (Figures 2N and 2N’) and thus expose higher PS. Indeed, we observed dendrite branches that traversed single *Eato*-KD cells (Figures 6K–6K”). Interestingly, Drpr was recruited to the sites of the dendrites on WT but not *Eato*-KD epidermal cells. Assuming that the level of PS exposure along the branch is relatively even, these results suggest that epidermal Eato acts upstream of Drpr recruitment.

Next, to test if supplying more Drpr can compensate for the loss of *Eato* in epidermal cells, we overexpressed Drpr in *en*+ epidermal cells of whole-body *Eato* KO animals. The *en*+ domain showed increased debris levels and reduced dendrites as compared to the anterior control region (Figures 6L– 6N), indicating rescue of engulfment. Thus, Eato sensitizes phagocytes by facilitating Drpr recruitment, but more Drpr can compensate for the reduction of sensitivity caused by *Eato* deficiency.

### Eato promotes engulfment activity of epidermal cells by suppressing PS exposure

Since Eato protects neurons by suppressing PS exposure on the cell surface, we wondered if Eato also inhibits PS exposure on phagocytes. Thus, we knocked out *Eato* using *tub-Cas9* (Poe et al., 2019)and examined PS exposure on epidermal cells using GFP-Lact. We observed a moderate (3.39 folds) increase in GFP-Lact labeling as compared to the WT control (Figures 7A–7C). Next, we asked whether *Eato* can suppress ectopic PS exposure on epidermal cells caused by flippase ablation. *CDC50* KO in epidermal cells resulted in strong labeling of GFP-Lact on the KO cells (Figure 7D), consistent with CDC50/ATP8A being the primary PS flippase on the plasma membrane (Sapar et al., 2018)(Sapar et al., 2018). Interestingly, overexpressing Eato in *CDC50* KO epidermal cells reduced Lact-GFP labeling to 34.3% of the original level (Figures 7E and 7F), confirming that Eato is capable of reducing PS exposure on epidermal cell surfaces. Next, to test if PS exposure contributes to the reduced phagocytosis of *Eato*-deficient epidermal cells, we co-expressed ATP8A and its chaperone CDC50 in *en*+ epidermal cells of whole-body *Eato* KO animals to suppress PS exposure. Indeed, the *en*+ domain showed elevated debris levels (Figures 7G and 7H), suggesting partial restoration of engulfment activity.

**Figure 7.**
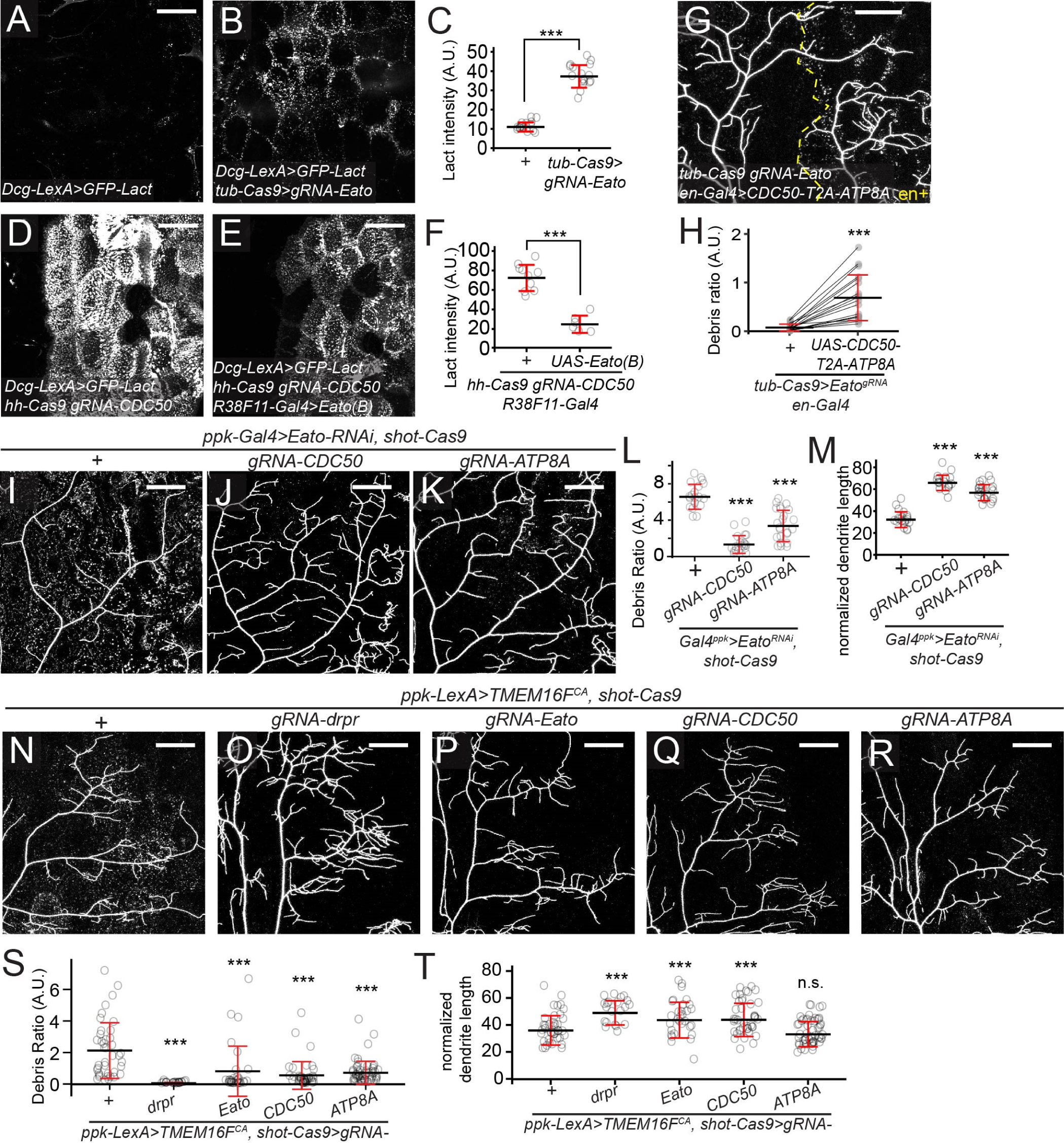
*Eato* promotes engulfment activity of epidermal cells by suppressing PS exposure. (A–C) Binding of the PS sensor GFP-Lact on control (A) and *Eato* KO (B) epidermal cells. *tub-Cas9* induces *Eato* KO ubiquitously. *dcg-Gal4* drives GFP-Lact expression in the fat body. Average fluorescence intensity of GFP-Lact on epidermal cells is quantified in (C). n = number of neurons and N = number of animals: *+* (n = 16, N = 8); *tub-Cas9>gRNA-Eato* (n = 17, N = 11). (D–F) GFP-Lact binding on *CDC50* KO (D) and *CDC50* KO with *Eato* overexpression (E) epidermal cells. *hh-Cas9* induces KO in the posterior half of each larval segment. *dcg-Gal4* drives GFP-Lact expression in the fat body. Images were acquired using a lower brightness setting than in (A) and (B) to avoid oversaturation. Average fluorescence intensity of GFP-Lact on *hh*+ epidermal cells is quantified in (F). n = number of neurons and N = number of animals: *hh-Cas9 gRNA-CDC50 R38F11-Gal4* (n = 10, N = 5); *hh-Cas9 gRNA-CDC50 R38F11-Gal4 UAS-Eato(B)* (n = 6, N = 4). (G–H) Dendrites of C4da neurons in whole-body *Eato* KO with CDC50-2A-ATP8A overexpression in *en*+ epidermal cells (G). The *en*+ domain is right to the yellow dashed line. Debris ratio in control (+) and CDC50-2A-ATP8A overexpressing domains is quantified in (H). Data is from 19 neurons in 12 animals. (I–M) Dendrites of C4da neurons with C4da-specific *Eato* KD (I) and additional epidermal *CDC50* KO (J), or additional epidermal *ATP8A* KO (K). Debris ratio is quantified in (L) and normalized dendrite length is quantified in (M). n = number of neurons and N = number of animals: *Gal4^ppk^>Eato^RNAi^ shot-Cas9* (n = 19, N = 13); *Gal4^ppk^>Eato^RNAi^ shot-Cas9 gRNA-CDC50* (n = 19, N = 13); *Gal4^ppk^>Eato^RNAi^ shot-Cas9 gRNA-ATP8A* (n = 23, N = 14). (N–T) Dendrites of C4da neurons with *TMEM16F^CA^* OE (N) and additional epidermal *drpr* KO (O), or additional epidermal *Eato* KO (P), or additional epidermal *CDC50* KO (Q), or additional epidermal *ATP8A* KO (R). Debris ratio is quantified in (S), and normalized dendrite length is quantified in (T). n = number of neurons and N = number of animals: *ppk-LexA>TMEM16F^CA^ shot-Cas9* (n = 40, N = 19); *ppk-LexA>TMEM16F^CA^ shot-Cas9 gRNA-drpr* (n = 21, N = 15); *ppk-LexA>TMEM16F^CA^ shot-Cas9 gRNA-Eato* (n = 30, N = 15), *ppk-LexA>TMEM16F^CA^ shot-Cas9 gRNA-CDC50* (n = 41, N = 25), *ppk-LexA>TMEM16F^CA^ shot-Cas9 gRNA-ATP8A* (n = 54, N = 24). C4da neurons were labeled by *ppk-MApHS* in (G), (I–K) and (N–R). Scale bars: 50 μm. In all plots, ***p<0.001; n.s., not significant. (C) and (F), t-test; (H), paired t-test; (L–M) and (S–T), one-way ANOVA with Tukey post-hoc test.

The above data imply that PS exposure on phagocytes negatively impacts phagocytosis. To test this idea directly, we ectopically induced PS exposure on epidermal cells by flippase KO and assayed the ability of epidermal cells to engulf degenerating dendrites. We first examined degenerating dendrites caused by neuronal KD of *Eato* (Figure 7I). Strikingly, *CDC50* KO in epidermal cells nearly completely suppressed the dendrite degeneration (Figures 7J, 7L and 7M). *ATP8A* KO showed a similar, albeit milder, suppression of dendrite degeneration (Figures 7K–7M). Next, we examined dendrite degeneration caused by neuronal expression of *TMEM16F^CA^* (Figure 7N). Again, *CDC50* KO in epidermal cells suppressed this type of dendrite degeneration as effectively as *drpr* KO and *Eato* KO, while *ATP8A* KO caused a lightly milder suppression (Figures 7O–7T).

Together, the above results suggest that PS exposure on phagocytes inhibits sensing of PS on the engulfment target and that Eato promotes engulfment at least partially by suppressing PS exposure on phagocytes.

### Lipid accumulation in cells is unlikely to account for the effects of *Eato* deficiency

Eato is thought to function as a floppase to export excessive lipids from *Drosophila* photoreceptors (Moulton et al., 2021). In these neurons, ineffective clearance of lipid accumulation caused by oxidative stress speeds up neurodegeneration (Moulton et al., 2021). To investigate whether lipid accumulation also contributes to the defects of *Eato*-deficient da neurons and epidermal cells, we first overexpressed Brummer (Bmm), a *Drosophila* triglyceride lipase, in *Eato* KO neurons. Bmm overexpression was previously shown to be effective in suppressing photoreceptor degeneration caused by lipid accumulation (Moulton et al., 2021). However, we did not observe any rescue of dendrite degeneration by Bmm overexpression (Figures S7A–S7D). Next, we used LSD-2-EGFP to visualize lipid droplets, which store excessive neutral lipids, in neurons and epidermal cells. However, we did not observe detectable increases of lipid droplets in either tissue upon whole-body *Eato* KO (Figures S7E–S7K). Although these results cannot completely exclude the possibility of lipid accumulation in *Eato*-deficient cells, they suggest that general lipid accumulation is not a significant cause of the defects of *Eato*-deficient cells.

### Human and *C. elegans* ABCA proteins are functional homologs of Eato

The human genome encodes 12 ABCA proteins involved in diverse biological processes (Albrecht and Viturro, 2007). The *C. elegans* CED-7 is a ABCA protein involved in phagocytosis (Ellis et al., 1991; Zhou et al., 2001). To explore potential functional conservation between Eato and these homologs, we tested if human and *C. elegans* ABCA genes can replace Eato’s functions in epidermal cells and neurons. Towards this aim, we obtained *UAS-hABCA1* and *UAS*-*hABCA2* from the Bloomington *Drosophila* Stock Center (BDSC) and generated *UAS*-driven hABC3, hABC4, hABC5, hABC12, and CED-7 in our laboratory. We also made a *UAS-mouse ABC7* (*mABC7*) transgene. In the following tests, *UAS-Eato(B)* served as a positive control while *UAS-Eato(B.MM)* served as a negative control; mammalian ABCA genes were expressed in flies at 29 ℃ to facilitate protein folding.

To test rescue of *Eato* LOF in epidermal cells, we expressed ABCA genes in the *en*+ domain of whole-body *Eato* KO animals. Interestingly, except mABCA7, all tested ABCA transgenes resulted in elevated levels of dendrite debris in the posterior hemi-segment (Figures 8A–8P), suggesting varying degrees of rescue. Among them, CED-7 rescued engulfment of dendrites as well as Eato(B), even though its effects on dendrite reduction and debris level are more variable (Figures 8D, 8O and 8P). The effects of human ABCA genes appear generally weaker than that of Eato(B) (Figures 8O and 8P). We next tested rescue of *Eato* LOF in neurons. However, none of the ABCA homologs obviously suppressed dendrite degeneration when expressed in *Eato*-KO C4da neurons, even though some of them resulted in reduced debris levels (Figures S8). These data suggest that most of the ABCA proteins examined share some similar biochemical activity that can boost engulfment activity of phagocytes, but they do not seem to function the same way as Eato in neurons.

**Figure 8.**
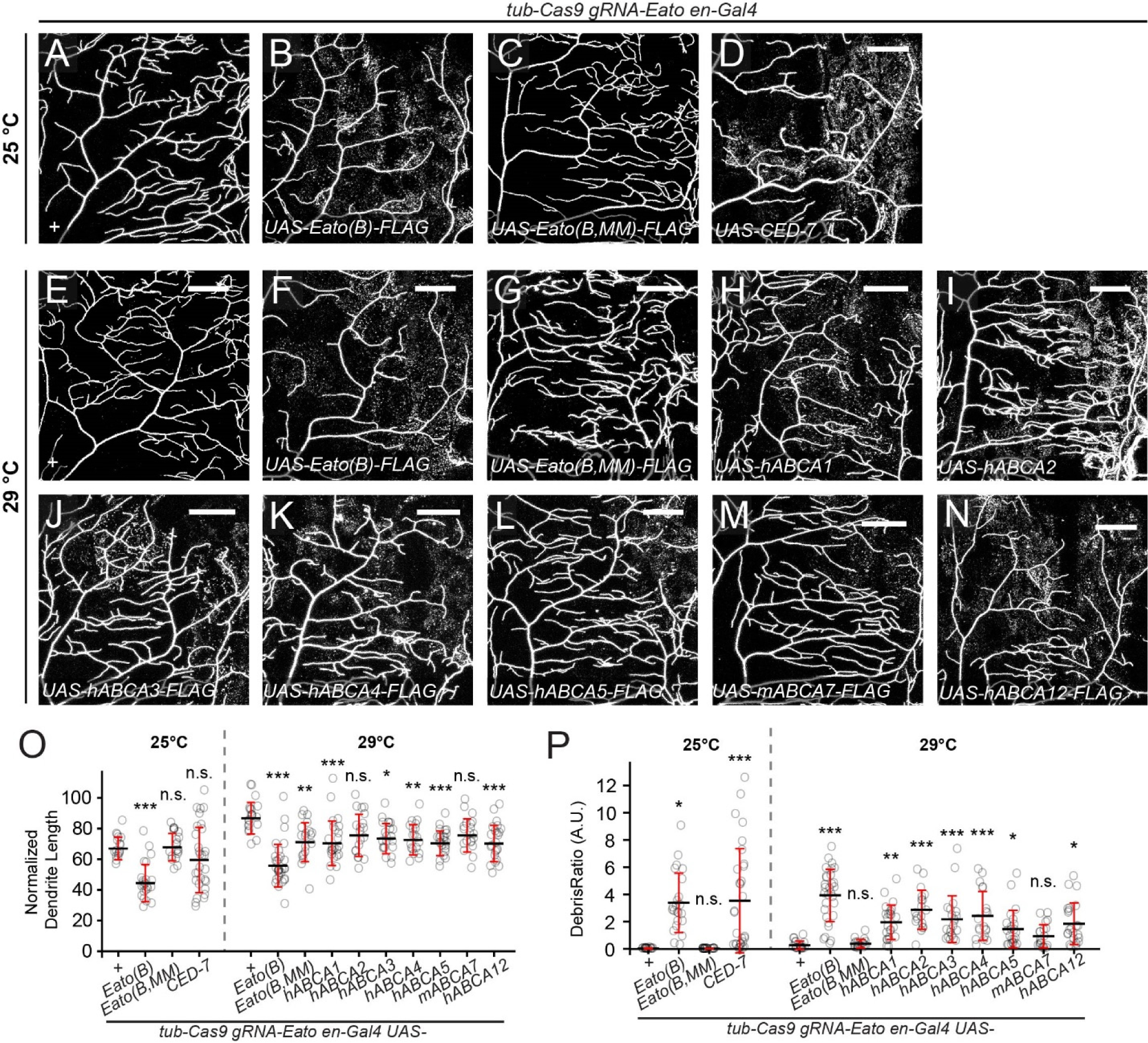
Human and *C. elegans* ABCA proteins are functional homologs of Eato. (A–D) Dendrites of C4da neurons in whole-body *Eato* KO (A) and with additional overexpression of *Eato(B)* (B), *Eato(B.MM)* (C), and *CED-7* (D) in the *en*+ domain. The larvae were raised at 25°C. (E–N) Dendrites of C4da neurons in whole-body *Eato* KO (E) and with additional overexpression of *Eato(B)* (F), *Eato(B.MM)* (G), human *ABCA1* (H), human *ABCA2* (I), human *ABCA3* (J), human *ABCA4* (K), human *ABCA5* (L), mouse *ABCA7* (M), and human *ABCA12* (N) in the *en*+ domain. The larvae were raised at 29°C. (O–P) Quantification of normalized dendrite length (O) and debris ratio (P) in A–N. n = number of neurons and N = number of animals. 25°C: + (n = 17, N = 10); *Eato(B)* (n = 23, N = 14); *Eato(B,MM)* (n = 20, N = 7); *CED-7* (n = 30, N = 11). 29°C: + (n = 20, N = 6); *Eato(B)* (n = 30, N = 13); *Eato(B,MM)* (n = 19, N = 6); *hABCA1* (n = 24, N = 7), *hABCA2* (n = 19, N = 6), *hABCA3* (n = 20, N = 6), *hABCA4* (n = 19, N = 6), *hABCA5* (n = 23, N = 5), *mABCA7* (n = 20, N = 6), *hABCA12* (n = 23, N = 7). Scale bar: 50 μm. In all plots, ***p<0.001; **p<0.01; *p<0.05; n.s., not significant; one-way ANOVA with Tukey post-hoc test.

## DISCUSSION

### Eato plays opposing roles in neurons and phagocytes to control phagocytosis-driven neurite degeneration

In this study, we discovered that a single ABCA transporter, Eato, plays opposite roles on the defensive and the offensive sides of phagocytosis-driven neurodegeneration. On the defensive side, Eato protects neurons from becoming targets of phagocytosis, but on the offensive side, Eato enhances the ability of phagocytes to detect nearby engulfment targets. Consequently, the loss of *Eato* in neurons alone causes surrounding phagocytes to attack and engulf the axons and dendrites, resulting in severe neurodegeneration. In contrast, removing *Eato* from both neurons and resident phagocytes prevents neurodegeneration because *Eato*-deficient phagocytes are no longer able to detect eat-me signals exposed on neurons. Interestingly, Eato’s opposing roles in neurons and phagocytes are both related to its ability to suppress PS exposure on the cell surface, suggesting a common biochemical activity underlying both phenotypes. Although multiple ABCA genes are known to be involved in neurodegeneration in model organisms or implicated in neurodegenerative human diseases, to our knowledge, such dual roles for ABCA genes in neurodegeneration have never been reported. Thus, Eato’s functions represent a new mechanism by which ABCA genes are involved in neurodegeneration.

Our results show that Eato protects diverse neurons in both the PNS and the CNS, suggesting that Eato is required for a general biological process shared by many neuronal types. However, *Eato* deficiency in some neuronal types (e.g. motor neurons) did not seem to cause degeneration (Figure S4H). At least two possibilities might explain this neuronal diversity. First, another ABCA gene may play similar roles as *Eato* in these neurons, such that the loss of *Eato* produces no effects. Along this line, we found that although Eato is broadly expressed in the nervous system, its expression is absent in some neurons. *Eato* LOF is not expected to cause degeneration of those neurons. Second, different neurons may interact with surrounding cells of different phagocytic capabilities, such that *Eato* mutant neurons are not engulfed if the neighboring cells are poor phagocytes. Supporting the idea of uneven phagocyte capacities, we found that epidermal cells are potent phagocytes that can eat most dendrites of live da neurons, while CNS glia cause only mild axon degeneration of da neurons on their own (Figure 3).

Several ABCA genes, including *Eato*, are known to promote phagocytosis (Hamon et al., 2000; Iwamoto et al., 2006; Santoso et al., 2018; Wu and Horvitz, 1998). Specifically, *Eato* is required for follicle cells to engulf dying nurse cells in the female germline (Santoso et al., 2018). Consistent with this finding, we found that *Eato* is also required for epidermal cells and glia to engulf dendrites and axons, respectively, of *Eato*-deficient da neurons. Considering that loss of neuronal *Eato* causes degeneration of diverse neurons while whole-animal *Eato* mutants show no signs of neurodegeneration, *Eato* must be widely required for phagocytes in the nervous system to engulf *Eato* mutant neurons. It was not previously known how Eato promotes phagocytosis. Using multiple models of dendrite degeneration, we show here that Eato is not required for engulfment per se; rather, it boosts the sensitivity of phagocytes towards PS-exposing targets. Eato does so by allowing for recruitment of the engulfment receptor Drpr to the site of engulfment, similar to the role of *C. elegans* CED-7 in recruitment of the Drpr homolog Ced-1 (Venegas and Zhou, 2007).

### PS exposure is responsible for the defects of *Eato* deficiency in both neurons and phagocytes

Mechanistically, our results support the idea that Eato exerts its function in both neurons and phagocytes by suppressing PS exposure. In both tissues, overexpression of the PS-specific flippase ATP8A can rescue the defects caused by the loss of *Eato*, suggesting that surface PS exposure is necessary for these defects. Meanwhile, Eato overexpression suppresses dendrite loss caused by ectopic PS exposure in neurons and dramatically reduces PS exposure induced by flippase KO in epidermal cells, suggesting that Eato can reduce cell surface PS exposure. Thus, it appears that Eato carries out similar biochemical activities in both cell types, resulting in less cell surface PS.

How does Eato suppress PS exposure? Previous work has linked two ABCA genes with both PS exposure and phagocytosis. Murine *ABCA1* is required for efficient clearance of apoptotic cells in the developing limb bud and for the phagocytic activity of macrophages (Hamon et al., 2000). Meanwhile, ABCA1 promotes Ca^2+^-induced PS exposure in blood cells (Hamon et al., 2000). In *C. elegans* embryos, *CED-7* is also required for efficient PS exposure on apoptotic cells (Venegas and Zhou, 2007). These observations are consistent with ABCA1 and CED-7 being lipid floppases that export lipids from the interior of cells. In contrast, the effect of Eato on PS exposure appears to be opposite to those of ABCA1 and CED-7 and is more similar to that of the flippase ATP8A. However, our results also show important distinctions between Eato and the PS flippase. On the one hand, flippase KO in epidermal cells results in high levels of PS exposure, whereas *Eato* KO produces a milder effect. On the other hand, *Eato* KO in neurons causes much more severe dendrite degeneration than flippase KO (Sapar et al., 2018). Thus, unlike P4-ATPases that import PS across the lipid bilayer and maintain general PS asymmetry on the plasma membrane, Eato seems to specifically suppress certain potent eat-me signals related to PS. One possibility to reconcile these observations is that Eato may function to selectively clear a subset of PS lipids that are particularly potent in inducing phagocytosis from the cell surface. *In vitro* analysis of Eato’s biochemical activity will be critical to establish how Eato regulates PS homeostasis at the plasma membrane.

How does PS exposure on phagocytes inhibit engulfment of PS-exposing dendrites? A simple hypothesis is that PS on the surface of phagocytes interacts with the engulfment receptor Drpr on the same membrane and thus interferes with Drpr’s ability to interact with PS exposed on dendrites. This hypothesis predicts that increasing the PS level on dendrites may outcompete PS on phagocytes and restore engulfment. Indeed, injury induces rapid and high PS exposure on severed dendrites (Ji et al., 2022; Sapar et al., 2018) and these dendrites can still be engulfed by *Eato*-KO epidermal cells.

Eato was previously linked to neurodegeneration through its role in exporting excessive lipids from photoreceptors (Moulton et al., 2021). In these cells, the loss of *Eato* results in accumulation of oxidized lipids inside lipid droplets, causing photoreceptors to die earlier in the presence of oxidative stress. However, we do not think a similar mechanism accounts for the neurodegeneration observed here. First, we did not detect lipid droplet increases in *Eato* mutant neurons. Second, reducing the lipid load inside neurons by lipase overexpression did not affect the degeneration of *Eato* mutant neurons.

Third, the *Eato* mutant neurons we examined here degenerate much more rapidly (within 5 days) than *Eato*-deficient photoreceptors exposed to oxidative stress (>20 days). Lastly, in the retina, glial cells protect photoreceptors by taking up excessive lipids from photoreceptors rather than being responsible for neurodegeneration by engulfing photoreceptors (Liu et al., 2015). Thus, we posit that, outside the retina, Eato plays a much broader protective role in the nervous system by suppressing PS exposure.

### Conserved and diverged functions of ABCA proteins

Besides human ABCA1 and CED-7, we found that several other ABCA proteins (hABCA2, hABCA3, hABCA4, hABCA5, and hABCA12) can rescue the phagocytic defects of *Eato* KO epidermal cells to various extents. These results are surprising, considering that ABCA proteins do not necessarily have the same biochemical activities (Quazi and Molday, 2011). For example, although most of the ABCA proteins characterized so far are involved in exporting lipids, ABCA4 is a flippase for N-retinylidene-phosphatidylethanolamine (Quazi et al., 2012). However, our results suggest that many, if not all, ABCA proteins may have some shared biochemical properties that can enhance phagocytosis.

Considering that Eato’s roles in neurons and phagocytes both involve suppression of PS exposure, it is further surprising that, although several ABCA proteins can compensate for the loss of *Eato* in epidermal cells, none of them can rescue *Eato* mutant neurons. One possibility to account for this difference is that Eato may require additional, neuronal-specific factors to function properly in neurons, and these factors do not interact with ABCA proteins derived from humans and worms. Another possibility is that neurons require more complete suppression of PS exposure than phagocytes to inhibit the effects of *Eato* LOF.

### Axons and dendrites contribute differently to overall neuronal integrity

An interesting finding from our results is that phagocytic damage to dendrites affects the integrity of axons much more than the other way around: Blocking engulfment of dendrites largely rescued axon degeneration of *Eato* KO neurons, but suppressing axon engulfment had little effect on dendrite degeneration. These results suggest that dendrites contribute more to the overall health of neurons than axons. One possible explanation is that axons may be more separated metabolically or spatially from the cell body than dendrites through cellular compartmentalization, (Glock et al., 2021; Overly et al., 1996) such that injury signals initiated in axons do not spread effectively to the cell body. Alternatively, dendrites are more important to the overall health of da neurons because they occupy a larger cellular volume than axons.

### Potential roles of ABCA genes in neurodegenerative diseases by regulating PS-mediated phagocytosis

Neuron-phagocyte interactions play important roles in the progress of neurodegeneration (Butler et al., 2021). Besides clearing dead neurons and debris of neurites, phagocytosis can promote or even drive neurodegeneration (Butler et al., 2021; Sapar and Han, 2019). For example, mutations in *Atp8a2* result in spontaneous axon degeneration and paralysis in mice, most likely due to phagocytosis of axons induced by ectopic PS exposure (Zhu et al., 2012). We recently found that disruption of NAD^+^ metabolism, which is common in neurodegenerative diseases (Fang et al., 2017; Verdin, 2015), can cause neurons to lose neurites due to PS-induced phagocytosis (Ji et al., 2022). Here we present another example where dysregulation of PS homeostasis on the plasma membrane results in phagocytosis-dependent neurodegeneration. Several human ABCA genes are associated with neurodegenerative diseases, including ABCA1, ABCA2, ABCA7 in Alzheimer’s disease, ABCA5 in Parkinson’s disease, and ABCA4 in macular degeneration (Bossaerts et al., 2022; Fu et al., 2015; Kim and Halliday, 2012; Piehler et al., 2012). It is an intriguing question whether any human ABCA protein is neuroprotective by suppressing PS exposure on neurons, like Eato in *Drosophila*. However, to address this question, it is important to investigate neuronal-specific LOF of ABCA genes in *in vivo* mammalian models, since potential neurodegeneration could be phagocytosis-dependent and the LOF of ABCA genes in phagocytes could suppress neurodegeneration.

## MATERIALS AND METHODS

### Drosophila strains

The fly strains used in this study are listed in Table S1 (Key Resource Table). In general, C4da neurons were labeled by *ppk-MApHS*, *ppk-CD4-tdTom*, or *ppk-Gal4 UAS-CD4-tdTom*; PS exposure on cell surface was visualized by *dcg-Gal4 UAS-GFP-Lact* or *dcg-LexA LexAop-GFP-Lact*.

Molecular cloning and transgenic flies, generation of *Eato* KI, Gal4, and mutant flies, CRISPR-TRiM, mosaic analysis, live imaging, injury, assay, dissection and staining, image analysis and quantification, statistical analysis are described in Supplemental methods. Plasmids are available from Addgene or upon request. *Drosophila* strains are available from Bloomington *Drosophila* Stock Center or upon request.

## Supporting information

Key resource table

## ACKNOWLEDGMENTS

We thank Huanghe Yang (Duke University), Rando Allikmets (Columbia University), Ding Xue (University of Colorado Boulder), Yannick Hamon and Dr. Giovanna Chimini (Aix Marseille University), and *Drosophila* Genomics Resource Center (DGRC) for plasmids; Marc Freeman (Vollum Institute) and Developmental Studies Hybridoma Bank for antibodies; Ben White (National Institute of Mental Health), Michael Welte (University of Rochester), and Bloomington *Drosophila* Stock Center for fly stocks; Cornell BRC Imaging facility for access to microscopes (funded by NIH grant S10OD018516); Cornell CSCU for advice on statistics; Michael Goldberg, Jeremy Baskin, Quan Yuan, and members of the Han lab for feedback on the manuscript. This work was supported by NIH grants (R01NS099125 and R24OD031953) awarded to C.H..

## AUTHOR CONTRIBUTIONS

Conceptualization: X.C., C.H.; Methodology: X.C., C.H.; Software: X.C., C.H.; Validation: X.C.; A.S., N.VR., A.Y.; Formal analysis: X.C.; N.VR., A.Y., R.C.; Investigation: X.C.; A.S., Z.H., N.VR., A.Y.; Resources: B.W.; Writing - Original Draft: X.C., C.H.; Writing - Review & Editing: X.C., C.H.; N.VR., A.Y.; Visualization: X.C.; Supervision: C.H.; Funding acquisition: C.H.

---

**Figure S1.**
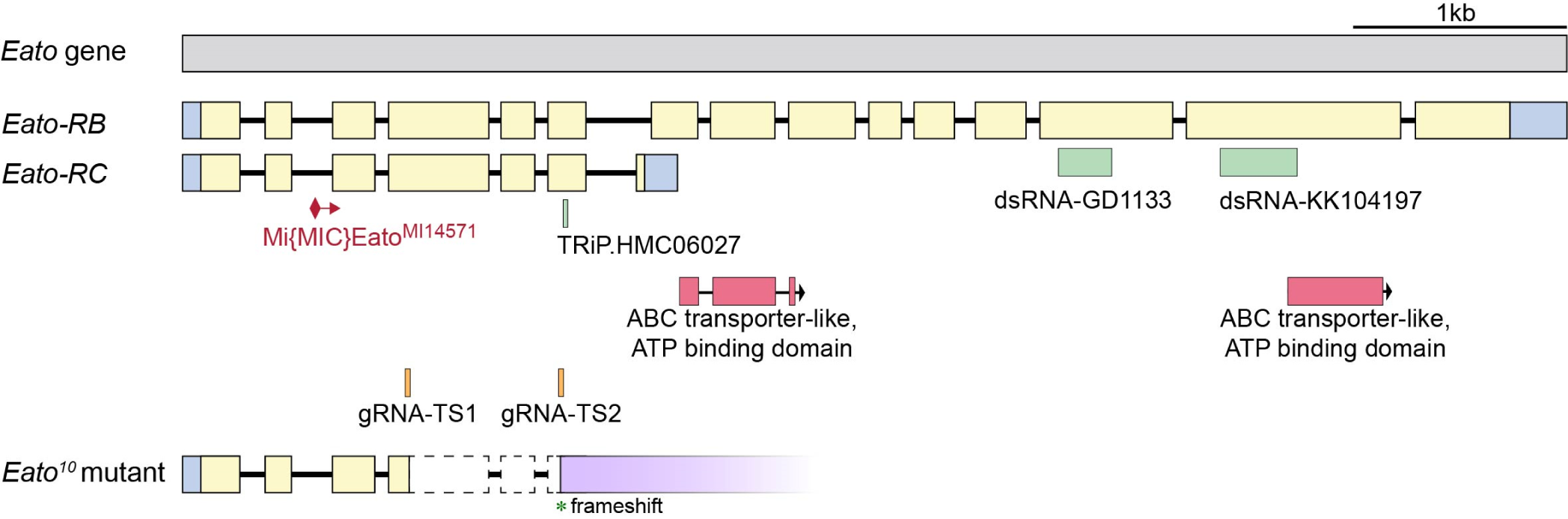
The gene structure of *Eato* and relevant reagents. *Eato* has two mRNA isoforms: *Eato-RB* (long) and *Eato-RC* (short). *Eato^MI14571^* (BDSC 59537) is a MiMIC insertion in the shared second intron of both isoforms. The RNAi line TRiP.HMC06027 (BDSC 65080) targets both isoforms whereas dsRNA transgenes GD1133 (VDRC) and KK104197(VDRC) target only the RB isoform. The locations of the two ATP-transporter-like ATP-binding domains are indicated. *gRNA-Eato* expresses two gRNAs targeting the shared sequence between RB and RC isoforms. *Eato^10^* contains a deletion from Exon 4 to Exon 6, causing frameshift after the deletion.

**Figure S2.**
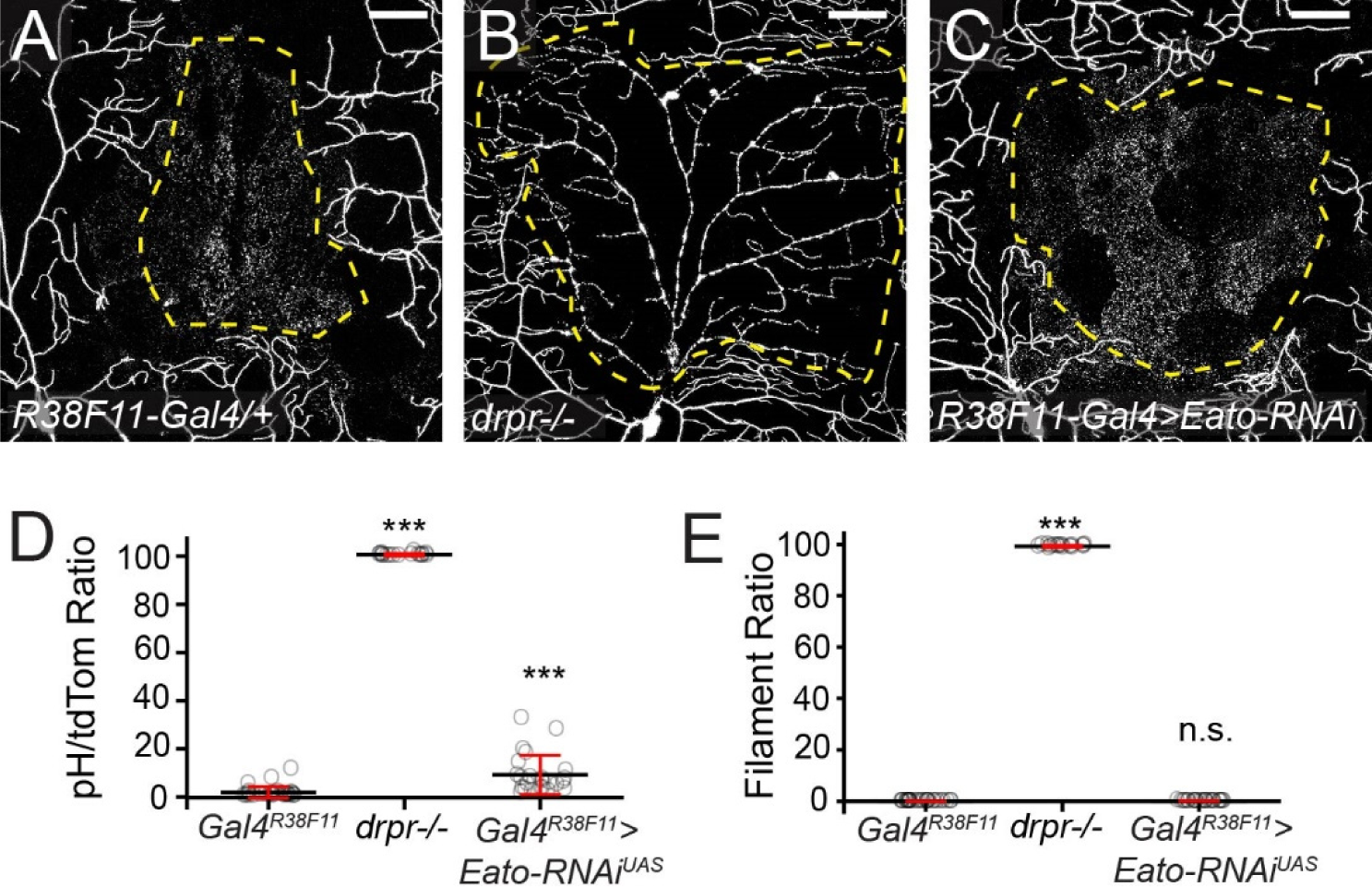
*Eato* LOF in epidermal cells does not block injury-induced degeneration. (A–C) C4da neuron dendrites in control (A), *drpr^indel3^* homozygous mutant (B), and epidermal cell-specific *Eato* KD (C) late 3^rd^ instar larvae 20h after laser injury. The dendrite territory before injury is enclosed by yellow dashed line. Scale bar: 50 μm. (D–E) Quantification of pHluorin/tdTomato ratio (D), which indicates the percentage of unengulfed materials, and the ratio of filament-like dendrite debris (E) in J–K. n = number of neurons and N = number of animals: control (n = 28, N = 10); *drpr^-/-^* (n = 17, N = 7); *Gal4^R38F11^ > Eato-RNAi^UAS^* (n = 24, N = 9). Two-sample t-test, ***p<0.001.

**Figure S3.**
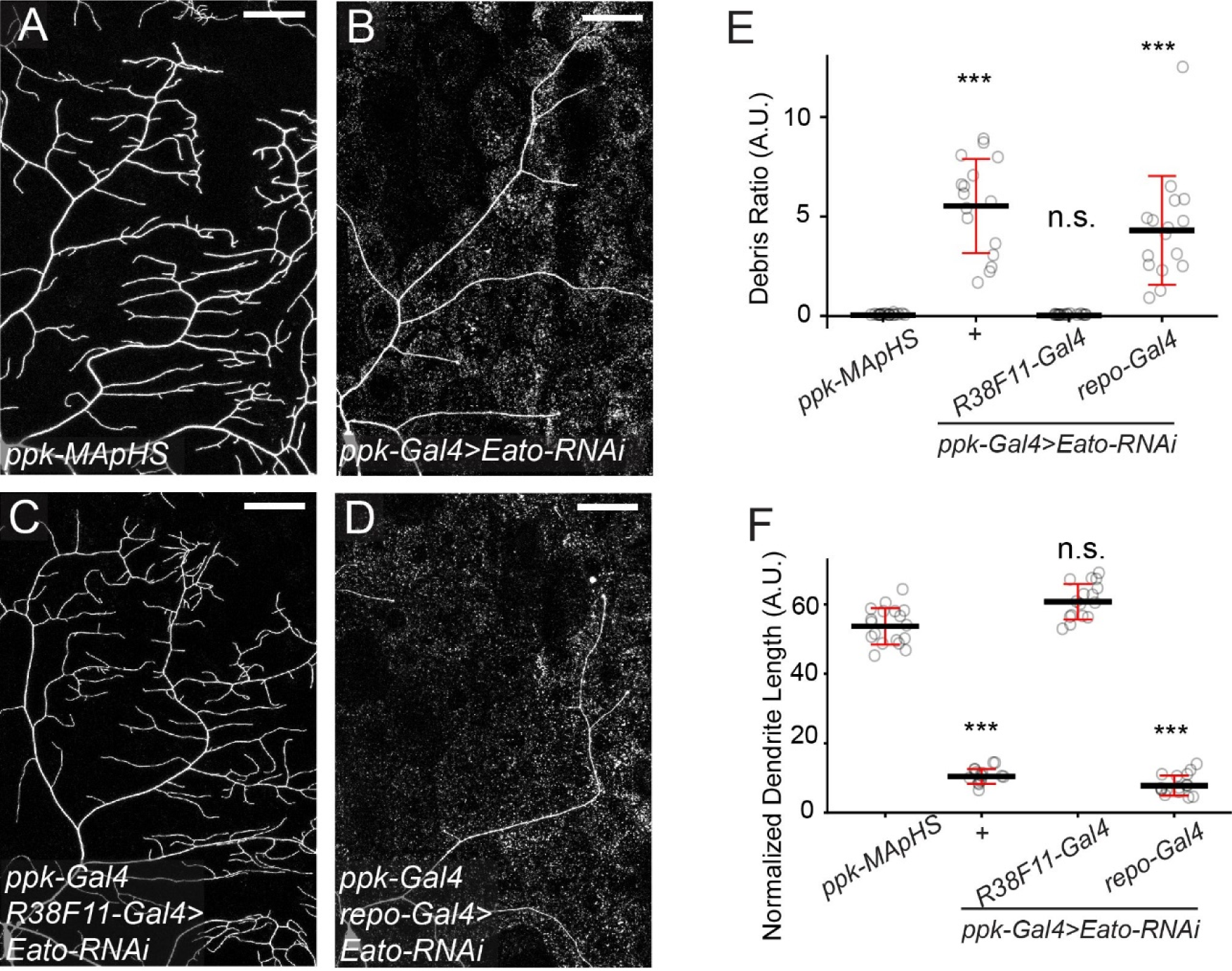
*Eato* LOF in glia does not block C4da dendrite degeneration. (A–D) Dendrites of C4da neurons in *ppk-Gal4* control (A), C4da-specific *Eato* KD (B), C4da and epidermal cell-specific *Eato* KD (C), and C4da and glia-specific *Eato* KD (D) late 3^rd^ instar larvae. Scale bar: 50 μm. (E–F) Quantification of debris ratio (E) and normalized dendrite length (F) in A–D. n = number of neurons and N = number of animals: *ppk-MApHS* (n = 18, N = 10); *ppk-Gal4>Eato-RNAi* (n = 16, N = 14); *ppk-Gal4 + repo-Gal4 >Eato-RNAi* (n = 16, N = 9), *ppk-Gal4 + R38F11-Gal4 >Eato-RNAi* (n = 16, N = 13). One-way ANOVA with Tukey post-hoc test, ***p<0.001.

**Figure S4.**
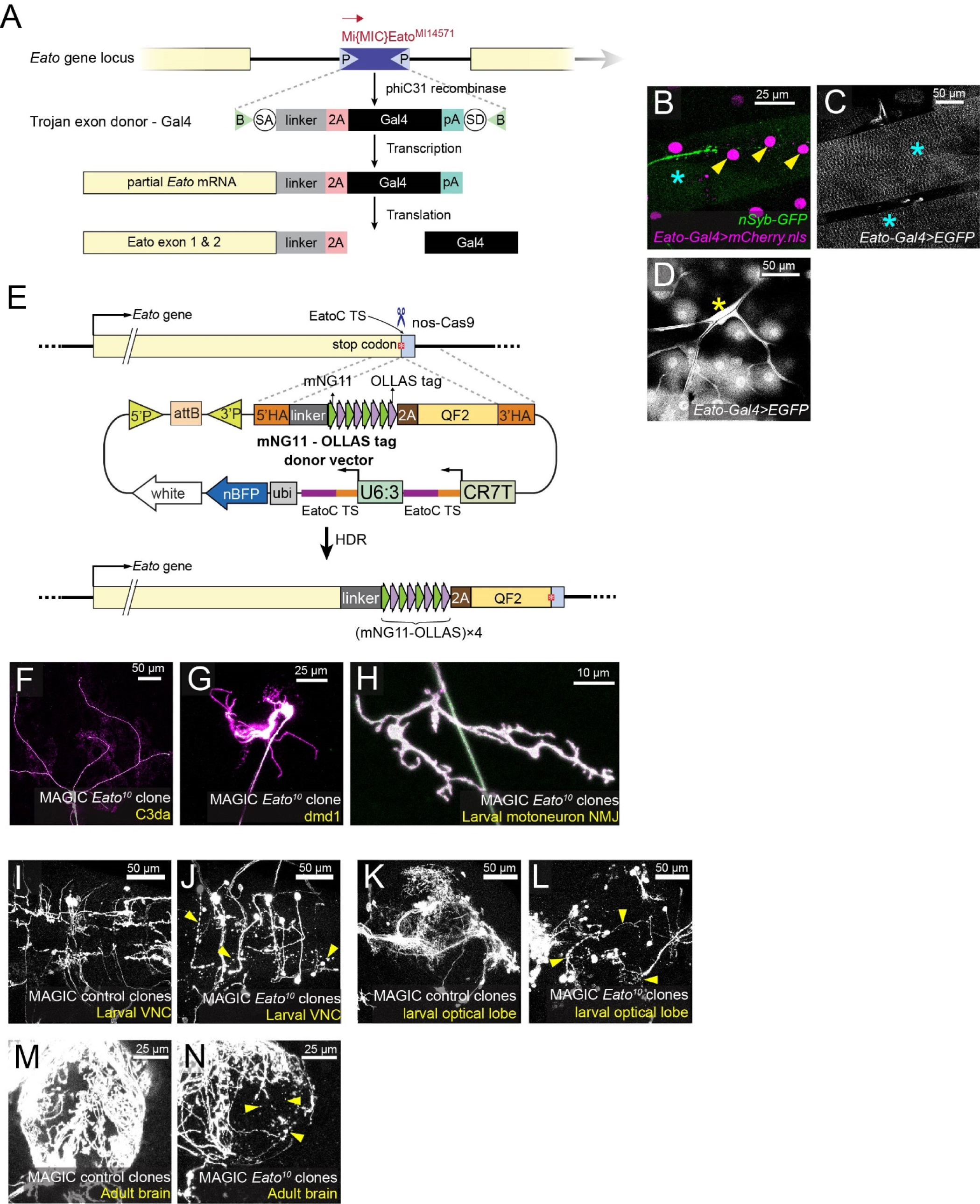
*Eato* LOF causes neuronal degeneration in both PNS and CNS. (A) A diagram showing the method of generating *Eato-Gal4* line. The 2A-Gal4 trojan exon is used to replace the MiMIC^MI14571^ insertion by recombinase-mediated cassette exchange. *T2A-Gal4* is expected to be spliced into the *Eato* mRNA, resulting in the expression of a truncated Eato protein and a Gal4 protein under the control of endogenous *Eato* regulatory sequence. (B–C) *Eato* expression in muscles. *Eato-Gal4^MI14571^* drives expression of a nuclear mCherry (B) and a cytosolic EGFP (C). Yellow arrowheads: muscle nuclei; cyan asterisks: muscle fibers. (D) *Eato* expression in tracheas. *Eato-Gal4^MI14571^* drives expression of a cytosolic EGFP. Yellow asterisk: tracheas. (E) A diagram showing the method of generating *Eato-(mNG_11_-OLLAS)_x4_* knock-in (KI) line. The gRNA-donor vector contains two gRNA expression units targeting the C-terminus of *Eato*. The (mNG_11_-OLLAS)_x4_-2A-QF2 cassette is flanked by 5’ and 3’ homology arms (HAs). Cas9-generated DNA break at the stop codon induces homologous recombination at the HAs. QF2 allows identification of KI lines by QUAS-driven reporters. A *ubi-nBFP* transgene on the vector allows negative selection of non-specific insertions of the vector in the genome. (F–G) *Eato^10^* mutant clones of Class III da (C3da) neuron (D) and dmd1 neuron (E) generated by MAGIC. (H) The neuromuscular junction of an *Eato^10^* mutant motoneuron clone. (I–N) Control (I, K, M) and *Eato* mutant (J, L, N) neuronal clones generated by MAGIC in the CNS, showing larval VNC (I–J), larval optical lobe (K–L), and adult brain (M–N). Yellow arrowheads indicate debris. Neurons were labeled by *RabX4-Gal4 UAS-MApHS* in (F–N). Both pHluorin (green) and tdTom (magenta) are shown in (F-H); only tdTom is shown in (I-N).

**Figure S5.**
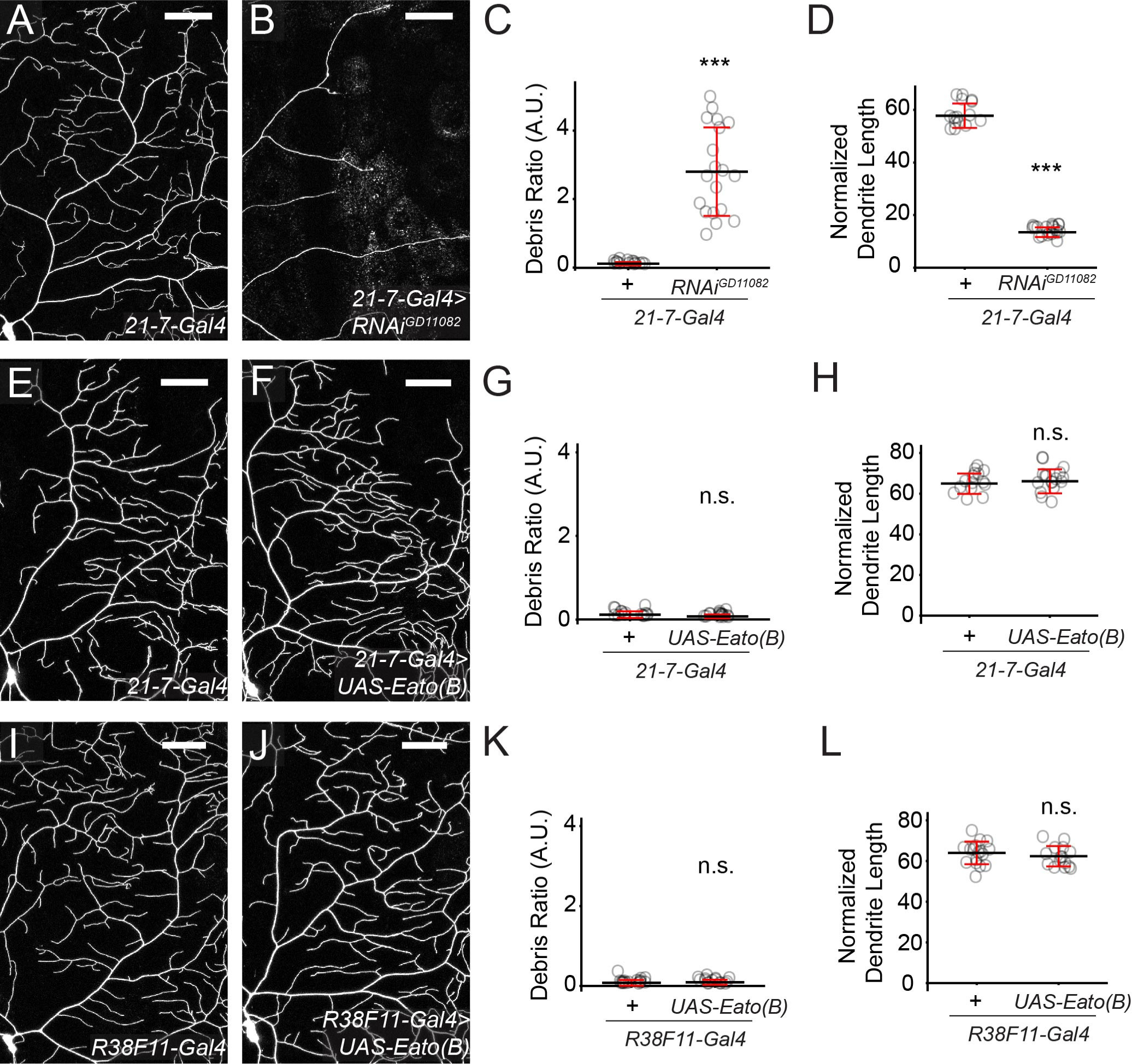
Eato overexpression in wildtype cells does not cause phenotypes. (A–D) Dendrites of C4da neurons in *21-7-Gal4* control (A) and da-specific *Eato(B)* KD (B) late 3^rd^ instar larvae. Debris ratio is quantified in (C) and normalized dendrite length is quantified in (D). n = number of neurons and N = number of animals: *21-7-Gal4* (n = 15, N = 8); *21-7-Gal4 RNAi^GD11082^* (n = 19, N = 11). (E–H) Dendrites of C4da neurons in *21-7-Gal4* control (E) and da-specific *Eato*(*B*) OE (F) late 3^rd^ instar larvae. Debris ratio is quantified in (G) and normalized dendrite length is quantified in (H). n = number of neurons and N = number of animals: *21-7-Gal4* (n = 15, N = 8); *21-7-Gal4 UAS-Eato(B)* (n = 17, N = 9). (I–L) Dendrites of C4da neurons in *R38F11-Gal4* control (I) and epidermal cell-specific *Eato(B)* OE (J) animals. Debris ratio is quantified in (K), and normalized dendrite length is quantified in (L). n = number of neurons and N = number of animals: *R38F11-Gal4* (n = 17, N = 9); *R38F11-Gal4 UAS-Eato(B)* (n = 15, N = 10). C4da neurons were labeled by *ppk-CD4-tdTomato* in (A–B); *ppk-MApHS* in (E–F) and (I–J). Scale bars: 50 μm. In all plots, ***p<0.001; n.s., not significant, two-sample t-test.

**Figure S6.**
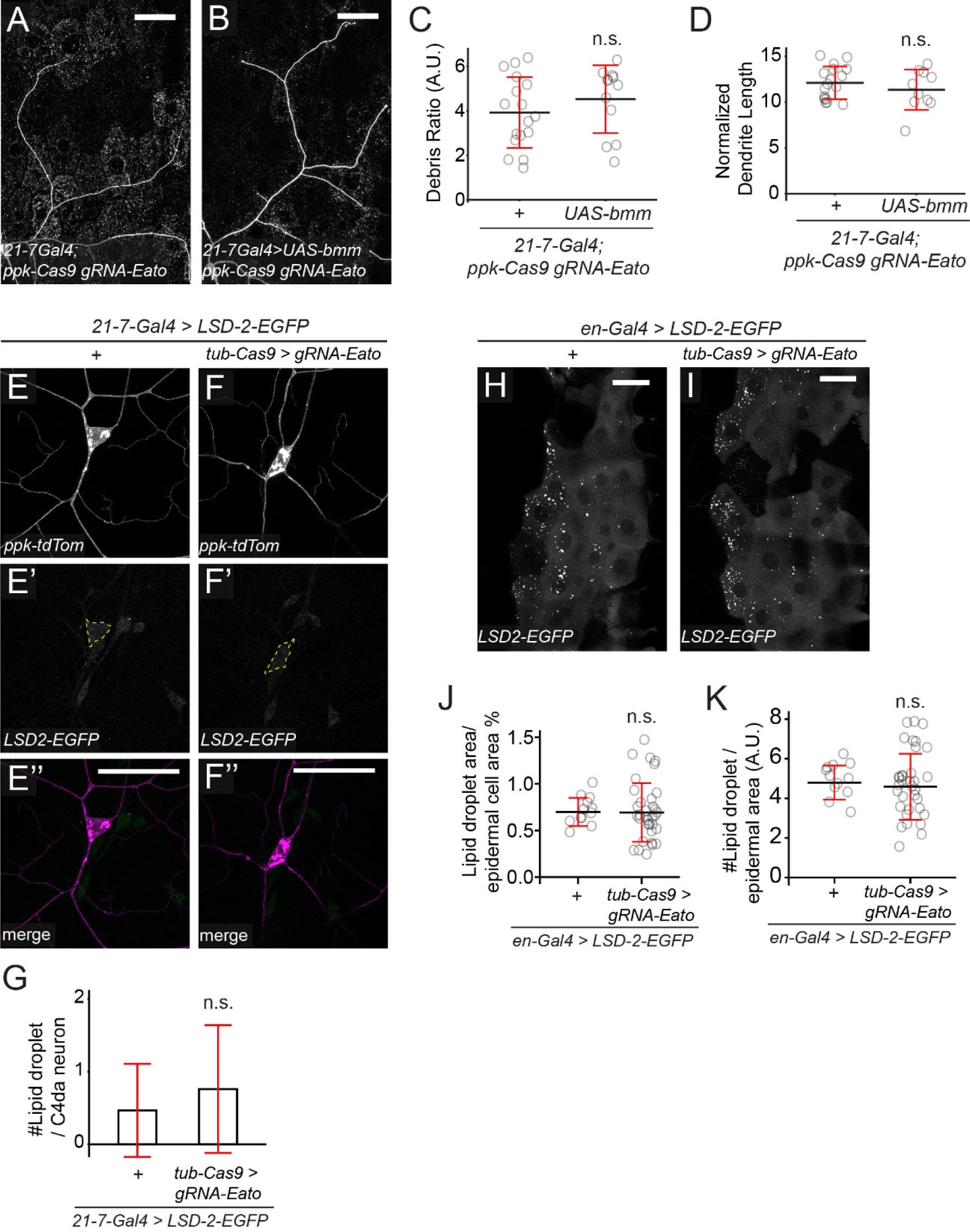
Lipid accumulation in cells does not seem to account for *Eato* deficiency. (A–B) Dendrites of *Eato* KO C4da neurons (A) and with additional overexpression of the lipase Bmm in da neurons (B). Debris ratio is quantified in (C), and normalized dendrite length is quantified in (D). n = number of neurons and N = number of animals: *21-7-Gal4 ppk-Cas9 gRNA-Eato* (n = 17, N = 11); *21-7-Gal4 UAS-bmm ppk-Cas9 gRNA-Eato* (n = 12, N = 6). (E–G) Distribution of the lipid droplet marker LSD-2-EGFP in control (E–E’’) and *Eato* KO (F–F’’) neurons. Number of LSD-2 puncta is quantified in (G). n = number of neurons and N = number of animals: *21-7-Gal4 > LSD-2-EGFP* (n = 15, N = 6); *21-7-Gal4 > LSD-2-EGFP tub-Cas9 gRNA-Eato* (n = 25, N = 9). (H–K) Distribution of the lipid droplet marker LSD-2-EGFP in control (H) and *Eato* KO (I) epidermal cells. Proportions of LSD-2 occupied area are quantified in (J) and number of LSD-2 puncta is quantified in (K). n = number of neurons and N = number of animals: *en-Gal4 > LSD-2-EGFP* (n = 12, N = 6); *en-Gal4 > LSD-2-EGFP tub-Cas9 gRNA-Eato* (n = 34, N = 11). Scale bar: 50 μm. In all plots, n.s., not significant; two-sample t-test.

**Figure S7.**
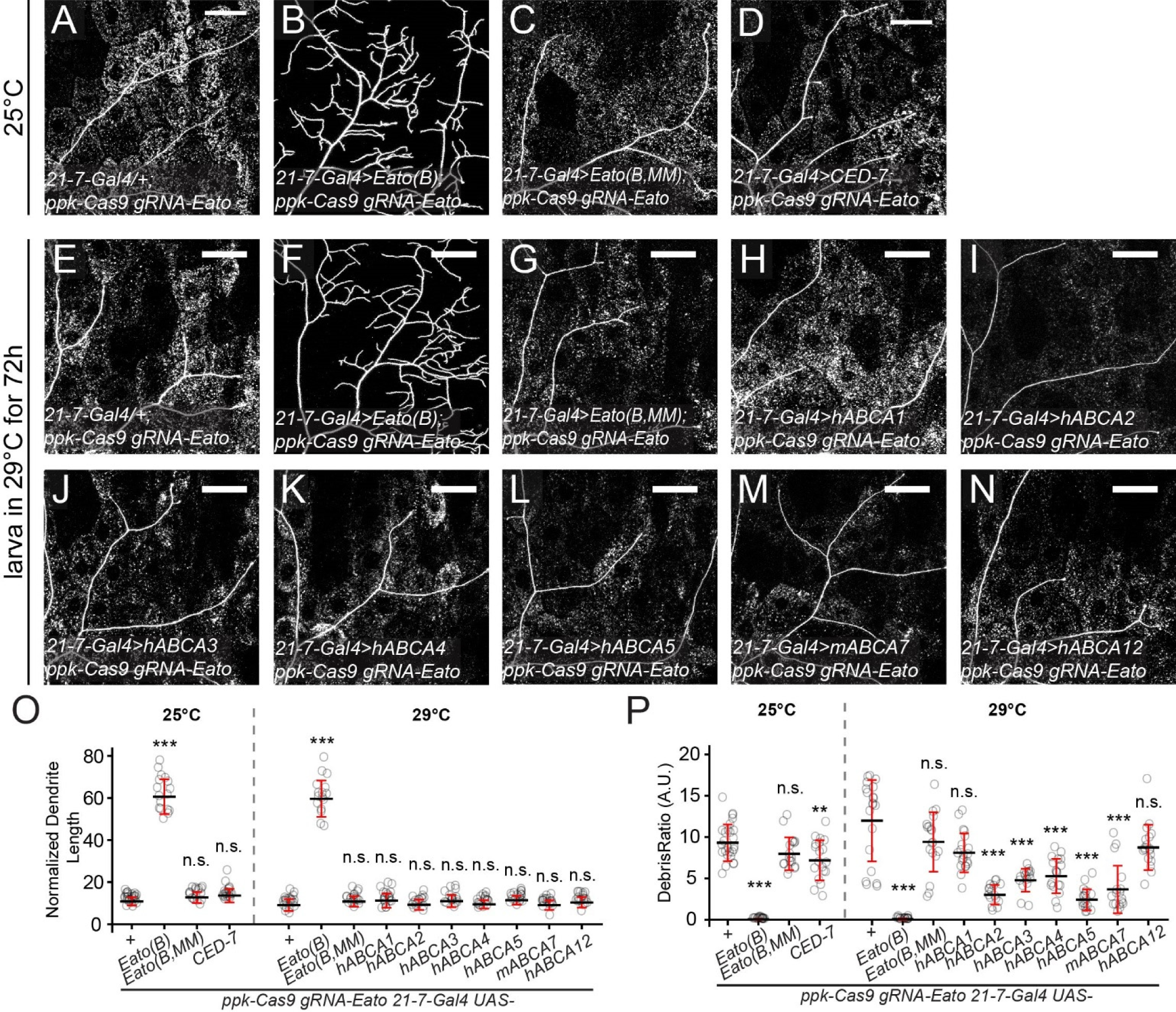
Human or *C. elegans* ABCA proteins cannot rescue *Eato* LOF in neurons. (A–D) Dendrites of *Eato* KO C4da neurons in control (A) and with da-specific overexpression of *Eato(B)* (B), *Eato(B.MM)* (C), and *CED-7* (D). The larvae were raised at 25°C. (E–N) Dendrites of *Eato* KO C4da neurons in control (A) and with da-specific overexpression of *Eato(B)* (F), *Eato(B.MM)* (G), human *ABCA1* (H), human *ABCA2* (I), human *ABCA3* (J), human *ABCA4* (K), human *ABCA5* (L), mouse *ABCA7* (M), and human *ABCA12* (N). The larvae were raised at 25°C for 24 hours and switched to 29°C for 72 hours. (O–P) Quantification of normalized dendrite length (O) and debris ratio (P) in A–N. n = number of neurons and N = number of animals. 25°C: + (n = 26, N = 13); *Eato(B)* (n = 17, N = 11); *Eato(B,MM)* (n = 18, N = 12); *CED-7* (n = 19, N = 8). 29°C: + (n = 20, N = 10); *Eato(B)* (n = 16, N = 8); *Eato(B,MM)* (n = 18, N = 9); *hABCA1* (n = 20, N = 10), *hABCA2* (n = 16, N = 8), *hABCA3* (n = 16, N = 8), *hABCA4* (n = 18, N = 9), *hABCA5* (n = 18, N = 9), *mABCA7* (n = 17, N = 8), *hABCA12* (n = 18, N = 8). *21-7-Gal4* drives expression of ABCA homologues in da neurons (A-N). Scale bar: 50 μm. In all plots, ***p<0.001; **p<0.01, n.s., not significant; one-way ANOVA with Tukey post-hoc test.

