## Supplemental methods for "Phagocytosis-driven neurodegeneration through opposing roles of an ABC transporter in neurons and phagocytes"

#### Molecular cloning and transgenic flies

*gRNA-Eato:* The gRNA expression vector was generated using published protocol (Koreman et al., 2021). The targeting sequences GCTCGGGAAAGACTTTGAGG and GTAGACCGAAAACATCGTTG were incorporated into cloning primers that were used to generate a PCR fragment with pTR(EF)-tRNA(Q) (Koreman et al., 2021) as the template. The PCR fragment was assembled with SapI-digested pAC-U63-tgRNA-Rev (Addgene 112811) (Poe et al., 2019) through NEBuilder HiFi DNA assembly (New England Biolabs, Inc) to make *Eato* gRNA expression vector.

*UAS-Eato(B)-FLAG: Eato(C)* coding sequence is PCR-amplified from BDGP clone FI17829 with oligos actctgaataGGGAATTGGGAATTcagaaaATGGCTGGGACTTTTAC and gttccttcacaaagatccTCTAGAttaCTTGTCGTCGTCGTCTTTGTAGTCGCTAGCATTTAACCCTGTTTGGTTATGCAGC. The fragment was cloned into EcoRI/XbaI digested pACU (Addgene 58372) (Han et al., 2011) by NEBuilder HiFi DNA assembly to make pACU-Eato(C)-FLAG. The UAS-Eato(C)-FLAG-SV40 fragment of pACU-Eato(C)-FLAG was PCR amplified using oligos gggcgaattggagctcgtttCGAAGAAAGGCCCACCCGTGAAGGTGAGCCctgcaggtcggagtactg and gggttttattaacttacatacatactagatccgatccagacatgataagatac and assembled into PmeI/EcoRI digested attB-P[acman]-ApR (DGRC 1245) (Venken et al., 2006) to make p[acman]US-Eato(C). The Eato(C) coding sequence was then removed by EcoRI/NheI digestion, and three overlapping DNA fragments (synthesized by Integrated DNA Technologies, Inc.) covering the whole coding sequence of *Eato(B)* were assembled into the EcoRI/NheI sites to make p[acman]US-Eato(B). The gRNA targeting sequences in Eato(B) were destroyed by silent mutations.

*UAS-Eato(B.MM)-FLAG*: Three overlapping DNA fragments (synthesized by Integrated DNA Technologies, Inc.) covering the whole coding sequence of *Eato(B)* were assembled into the EcoRI/NheI digested p[acman]US-Eato(C). The resulting construct p[acman]US-Eato(B.MM) contains K503M and K1293M mutations introduced into the synthetic DNA fragments.

*UAS-hABCA3-FLAG*: The *hABCA3* coding sequence was synthesized in three overlapping DNA fragments and assembled into EcoRI/NheI digested pACU-Eato(C)-FLAG to make pACU-hABCA3-FLAG.

*UAS-hABCA4-FLAG*: A new UAS vector pIACU3 was first constructed by replacing His2Av polyA of pIHEU with SV40 polyA and removing the synthetic intron after Hsp70 core promoter. Then a fragment containing AA2263-AA2273 of hABCA4 and a FLAG tag was introduced into pIACU3 between XhoI/XbaI sites. The remaining coding sequence of hABCA4 was isolated as two restriction fragments (NotI/XbaI and XbaI/XhoI) from pRK5-ABCA4 (a gift from Dr. Rando Allikmets, Columbia University) and ligated into the NotI/XhoI sites of pIACU3-FLAG to generate pIACU3-hABCA4-FLAG.

*UAS-hABCA5-FLAG*: The *hABCA5* coding sequence was synthesized in three overlapping DNA fragments and assembled into EcoRI/NheI digested pACU-Eato(C)-FLAG to make pACU-hABCA5-FLAG.

*UAS-mABCA7-YFP*: The *mABCA7-YFP* coding sequence was isolated from pBudCE4-ABCA7:YFP (a gift from Dr. Yannick Hamon and Dr. Giovanna Chimini, Aix Marseille University) by NotI/MluI digestion. The fragment was ligated with NotI/XbaI digested pACU and annealed oligos CGCGTaccggTTAATTAAggT and CTAGAccTTAATTAAccggtA to make pACU-ABCA7-YFP.

*UAS-hABCA12-FLAG*: The *hABCA12* coding sequence was synthesized in three overlapping DNA fragments and assembled into EcoRI/NheI digested pACU-Eato(C)-FLAG to make pACU-hABCA12-FLAG.

*UAS-CED-7-FLAG*: The plasmid pSL1190-Ced7 (a gift from Dr. Ding Xue, University of Colorado Boulder) was digested by NheI/SpeI to isolate *ced-7* coding sequence, which was then cloned into pACU. The resulting construct was then digested by AvrII/NotI and ligated with annealed oligos of ctaggGACTACAAAGACGACGACGACAAGtaagc and ggccgcttaCTTGTCGTCGTCGTCTTTGTAGTCc to introduce a FLAG tag at the C-terminus of CED-7.

*UAS-mNG(1-10)*: A synthetic DNA fragment encoding the 1-10 fragment of mNeonGreen2 (Feng et al., 2017) was assembled into EcoRI/XbaI sites of pIACU3 to generate pIACU3-mNG2(1-10).

*LexAop-TMEM16F(Y563K-D703R)*: A new LexAop2 construct, pAPLO3, was first constructed by removing the synthetic intron after Hsp70 core promotor in pAPLO (Poe et al., 2017). Two copies of LexAop2 were unintentionally mutated during the process, resulting in a 11xLexAop2 construct. The murine TMEM16F coding sequence containing Y563K and D703R mutations were PCR-amplified from pEGFP-N1-m16F(Y563K-D703R) (a gift from Dr. Huanghe Yang, Duke University) and ligated into EcoRI/XbaI sites of pAPLO3 to make pAPLO3-TMEM16F(Y563K-D703R).

*UAS-CDC50-T2A-ATP8A(core)*: A CDC50-T2A fragment was isolated from pACU-CDC50-T2A (Ji et al., 2023) by PacI/NheI digestion and then inserted into PacI/NheI sites of pACU-ATP8A(core) (Ji et al., 2022) to make pACU-CDC50-T2A-ATP8A(core).

*tub-Cas9*: An enhancer from *α-tub84B* was PCR-amplified from *w1118* genomic DNA using oligos ggggACAAGTTTGTACAAAAAAGCAGGCTGCTTGCACAGGTCCTGTTC and ggggACCACTTTGTACAAGAAAGCTGGGTATTCACGCTGTGGATGAGGAG and used to create an entry vector (pENTR221-tubP) through a Gateway BP reaction (Thermo Fisher Scientific). pENTR221-tubP was then combined with pDEST-APIC-Cas9 (Addgene 121657) (Poe et al., 2017) to generate pAPIC-tub-Cas9 expression vector through a Gateway LR reaction.

*ey-Cas9*: Three copies of 211 bp *ey* enhancer (corresponding to nucleotides 2577 to 2787 of GenBank accession number AJ131630) (Newsome et al., 2000) was first cloned into the XhoI/SalI sites of pENTR11 through a Gateway BP reaction (Thermo Fisher Scientific) to generate pENTR11-ey. pENTR11-ey was then combined with a Cas9 destination vector to generate the expression vector pAPIC-repo-Cas9. The Cas9 destination vector is similar to pDEST-APIC-Cas9 (Addgene 121657) but does not contain Inr, MTE, and DPE in the Hsp70 core promoter.

*Eato-(mNG*$_{11}$*-OLLAS)*$_{x4}$*-T2A-QF2* gRNA-donor vector: To make *Eato* knock-in (KI) flies, a combined gRNA-donor vector was constructed in pAC-CR7T-gRNA2.1-nlsBFP (Addgene 170515) (Koreman et al., 2021). First, two copies of the target sequence CTACGTTTAGGCCAGATAGT were introduced into SapI digested vector according to published protocols (Koreman et al., 2021), so that one gRNA is driven by CR7T promoter and another by U6:3 promoter. The resulting plasmid was digested by PstI and NheI and assembled with three DNA fragments to make the final gRNA-donor vector. The three DNA fragments include a (mNG$_{11}$-OLLAS)$_{X4}$-T2A-QF2 DNA fragment (synthesized by IDT, Inc.), and 5' and 3' homology arms (surrounding the stop codon of *Eato*, ~1 kb each) that were PCR-amplified from the genomic DNA of $w^{1118}$.

Transgenic constructs were injected by Rainbow Transgenic Flies to transform flies through φC31 integrase-mediated integration into attP docker sites.

### Generation of *Eato* KI flies

To generate *Eato-(mNG*$_{11}$*-OLLAS)*$_{x4}$*-T2A-QF2*, the *Eato* gRNA-donor vector was injected into *y1 nos-Cas9ZH-2A w** (BDSC, #54591) embryos. Adult flies from injected embryos were crossed to *10×QUAS-6×GFP/CyO,weeP* (BDSC#52264). The larval progeny was screened for GFP-positive and BFP-negative animals. Candidates were collected to cross with *Sco/CyO,weeP* individually to remove *10×QUAS-6×GFP* and *nos-Cas9ZH-2A* and to establish isogenic stocks. The *(mNG*$_{11}$*-OLLAS)*$_{x4}$*-T2A-QF2* insertion was verified by genomic PCR and sequencing.

**Generation of *Eato-Gal4* flies**

*Eato-Gal4* was converted from *Eato*$^{MiMIC[MI14571]}$ (BDSC #59537) line using Trojan Exon method (Diao et al., 2015). *Cre vas-int/Y*; *Eato*$^{MiMIC}$*/+; pC(loxP2-attB-SA(1)-T2A-Gal4-Hsp70)/+* males were crossed with *UAS-2×EGFP* females for a screen of GFP-positive larva. GFP-positive male progenies were crossed to *Sco/CyO, wee-P* females to remove other components except *Eato-Gal4*. The successful RMCE event was confirmed by genomic PCR and sequencing.

**Generation of *Eato* mutant by CRISPR/Cas9**

*gRNA-Eato* flies were crossed to *nos-Cas9*$^{attP2}$ flies. Female progenies were crossed to *Sp*/*CyO,weeP*; *Tm2*/*Tm6B*,Tb flies and individual progenies were collected and crossed to *Sp*/*CyO,weeP*; *Tm2*/*Tm6B*,Tb again for amplification. Single flies picked from the progeny was used in genomic PCR to screen for deletions between the two gRNA target sites. *Eato*$^{10}$ mutant line used in this study contains a deletion between exon 4 and 6, resulting in a 709bp deletion and reading frame shift from amino acid 201.

**CRISPR-TRiM**

The efficiency of *gRNA-Eato* was validated by the Cas9-LEThAL assay (Poe et al., 2019). Homozygous males of each gRNA line were crossed to *Act-Cas9 w lig4* homozygous females. *gRNA-Eato* crosses yielded viable female progeny and prepupal male lethality, suggesting that the gRNA is efficient.

Tissue-specific gene knockout in Class IV da neurons was done by using *ppk-Cas9* (Poe et al., 2019). Knockout in pan-epidermal cells was carried out using *shot-Cas9* (Ji et al., 2022). Knockout in posterior half of epidermal in each segment was carried out using *hh-Cas9* (Poe et al., 2019). Whole-animal knockout was carried out using *tub-Cas9*.

**Mosaic analysis by gRNA-induced crossing-over (MAGIC)**

The principle of MAGIC method was previously described (Allen et al., 2021). Briefly, gRNA-induced double strand breaks cause crossovers between homologous chromosomes in precursor cells, which can result in homozygous daughter cells after chromosomal segregation. A gRNA-marker transgene, which expresses ubiquitously Gal80 and gRNAs targeting a centromere-proximal region, is located on the chromosomal arm of interest (2L) and is paired with the mutant chromosome. The homozygous mutant clones generated by this way will lose Gal80 and thus can be labeled by Gal4-driven expression of a fluorescent marker. *gRNA-40D2(Gal80)* was used as the gRNA-marker for chromosomal arm 2L (Allen et al., 2021). Homozygous *Eato*$^{10}$ mutant clones were labeled by *RabX4-Gal4 UAS-MApHS*.

Clones in larval peripheral neurons were induced by *zk-Cas9*, which is expressed in precursor cells of peripheral neurons and epidermal cells (Allen et al., 2021). Clones were induced by *ey-Cas9* in the larval optical lobe, *SOP-Cas9* (Poe et al., 2019) in the larval ventral nerve cord, and *HS-Cas9* (Garcia-Marques et al., 2020) in the adult brain.

**Live imaging**

Live imaging was performed as previously described (Ji et al., 2022). Briefly, animals were reared at 25˚C in density-controlled vials (~100 embryos/vial) for 96 h, 120 h, or as specified in standard yeast-glucose medium (doi:10.1101/pdb.rec10907). Larvae were mounted in glycerol and imaged using a Leica SP8 confocal microscope. For consistency, only dorsal ddaC neurons of A2 and A3 segments (2 neurons per animal) on one side of the larvae were imaged.

**Injury assay**

Injury assay at the larval stage was done as described previously (Sapar et al., 2018). Briefly, larvae at 72 h AEL were lightly anesthetized with isoflurane and the primary dendrites of ddaC neurons in segment A2 and A3 were ablated using 790 nm two-photon laser on a Zeiss LSM880 Confocal/Multiphoton Upright Microscope. Animals were recovered on grape juice agar plates after ablation for 20 h before imaging.

**Larval brain preparation**

Larval brain dissection was performed as described previously (Belenkaya et al., 2004). Briefly, wandering 3rd instar larvae were dissected in a small petri dish filled with cold PBS. The anterior half of the larva was inverted, and the trachea and gut were removed. Samples were then transferred to 4% formaldehyde in PBS and fixed for 25 minutes at room temperature. Brain samples were washed with PBS. After immunostaining, the brains were placed in SlowFade Diamond Antifade Mountant (Thermo Fisher Scientific) on a glass slide. A coverslip was lightly pressed on top. Brains were imaged with 40× NA1.3 oil objective using a Leica SP8 confocal microscope.

**Larval fillet preparation**

Larval fillet dissection was performed on a petri dish half-filled with PMDS gel. Wandering third instar larvae were pinned on the dish in cold PBS, ventral-side up and then dissected to expand the body wall. PBS was then removed and 4% formaldehyde in PBS was added to fix larvae for 15 minutes at room temperature. Fillets were rinsed and then washed at room temperature in PBS for 20 minutes. After immunostaining, the head and tail of fillets were removed, and the remaining fillets were placed in SlowFade Diamond Antifade Mountant on a glass slide. A coverslip was lightly pressed on top. Larval fillets were imaged with 40× NA1.3 oil objective using a Leica SP8 confocal microscope.

**Immunohistochemistry**

For larval brains: Following fixation, brains were rinsed and then washed twice at room temperature in PBS with 0.3% Triton-X100 (PBST) for 20 minutes each. Brains were then blocked in a solution of 5% normal donkey serum (NDS) in PBST for 1 hour. Brains were then incubated in the blocking solution with rat mAb 7E8A10 anti-Elav (1:10 dilution, DSHB) or mouse mAb 8D12 anti-Repo (1:20 dilution, DSHB) overnight at 4°C. Following incubation, brains were then rinsed and washed in PBST 3 times for 20 minutes each. Brains were then incubated in a blocking solution containing a donkey anti-rat or donkey anti-mouse secondary antibody conjugated with Cy5 or Cy3 (1:400 dilution, Jackson ImmunoResearch) for 2 hours at room temperature. Brains were then rinsed and washed in PBST 3 times for 20 minutes each and stored at 4˚C until mounting and imaging.

For larval fillets: Following fixation, fillets were rinsed and then washed at room temperature in PBS. Fillets were then removed from PMDS gel and blocked in a solution of 5% normal donkey serum (NDS) in 0.2% PBST for 1 hour. Fillets were then incubated in the blocking solution with primary antibodies overnight at 4°C. Primary antibodies used in this study are V5 Tag Antibody (R960-25) (1:400 dilution, Thermo Fisher Scientific), Rat anti OLLAS Epitope Tag Antibody (L2) (1:100 dilution, Invitrogen), and Rabbit anti Drpr polyclonal antibody (1:100 dilution, a gift from Dr. Marc Freeman). Following incubation, fillets were then rinsed and washed in PBST 3 times for 20 minutes each. fillets were then incubated in a blocking solution containing fluorophore-conjugated conjugated secondary antibodies for 2 hours at room temperature. Secondary antibodies used in this study were: donkey anti-mouse secondary antibody conjugated with Cy5 or Alexa 488 (1:400, Jackson ImmunoResearch), donkey anti-rabbit secondary antibody conjugated with Cy5 or Alexa488 (1:400, Jackson ImmunoResearch) and donkey anti-Rat secondary antibody conjugated with Cy5 (1:400, Jackson ImmunoResearch). Fillets were then rinsed and washed in PBST 3 times for 20 minutes each and stored at 4˚C until mounting and imaging.

**Image analysis and quantification**

Dendrites morphology and debris formation: Image processing and analyses were done in Fiji/ImageJ. The method to measure the debris resulting from dendrite degeneration has been described(Sapar et al., 2018). Briefly, for MApHS-labelled neurons, pHluorin channel was segmented to generate a dendrite mask and tdTomato channel was segmented to include signals from both remaining dendrites and debris. The pHluorin area was subtracted from dendrites + debris area (tdTomato) to generate a mask containing only debris signal.

For tdTomato-labelled neurons, a fixed threshold was used to segment both dendrites and debris. The debris signal was later separated from dendrite signal based on different size and circularity. A region of interest (ROI) was manually drawn to include a quadrant of a ddaC neuron's territory. The methods for tracing and measuring C4da neuron dendrite length have been previously described (Poe et al., 2017). The ROI area, total dendrite length and debris area were measured. Dendrite length and debris area were normalized by ROI area for comparison between different genotypes.

The method to measure injured dendrites marked by ppk-MApHS was previously described (Ji et al., 2022). Briefly, the pHluorin-positive area was divided by tdTomato-positive area in an ROI to produce unengulfment ratio. For measuring debris dispersion of injured dendrites, dendrite debris was segmented by Auto Threshold (the "Default" method) in a rectangular ROI that was previously covered by injured dendrites. The ROI was divided into 15X15-pixel squares. The debris spread index was the area ratio of all squares containing dendrite debris in the ROI.

<u>Axon morphology</u>: VNC images are evaluated with a score system. A score of 0-3 was given to each segment with no obvious morphological change (0), debris formed (1), axon blebbing (2) and axon fragmentation (3). The degeneration score of one VNC is the sum of the scores of all 12 segments.

**Experimental design and statistical analysis**

For all experiments, the control groups and the experimental groups were kept in the same growing conditions. The same dissection and staining procedures were applied to all the groups. The animals used for dissection were of the same age. R was used to perform one-way analysis of variance (ANOVA) and t-test where indicated. Non-equal variance was assumed. For experiments with more than two groups, one-way ANOVA was first applied to identify significantly different mean(s). After that, multiple comparisons were performed using the Tukey post hoc method.
